# Pervasive translational control of photosynthesis genes during photomorphogenesis is acquired by C_4_ genes

**DOI:** 10.1101/2023.10.27.563924

**Authors:** Ivan Reyna-Llorens, Filip Lastovka, Tina B. Schreier, Pallavi Singh, Betty Y.W. Chung, Julian M. Hibberd

## Abstract

C_4_ photosynthesis allows increased efficiency and has evolved in more than sixty-six plant lineages. Underpinning this repeated appearance of the C_4_ pathway is a major transcriptional reprogramming of photosynthesis genes. Here we investigated whether evolution has also significantly modified translational control by defining the translational dynamics of C_3_ rice and C_4_ sorghum during photomorphogenesis. In the dark rice photosynthesis transcripts are low abundance but highly translated. After exposure to light translational efficiency declines. The same phenomena occur in sorghum but in addition C_4_ cycle genes show this response. We propose a model in which translational control of photosynthesis genes permits a rapid response to light and that this translational regulation is gained by C_4_ genes during the evolution of the C_4_ pathway.

## Introduction

After perceiving light etiolated leaves respond rapidly to become photosynthetic and in so doing maximize their metabolic potential. This is a highly conserved process that operates in C_3_ plants but also those that use the derived C_4_ pathway. To date the majority of downstream signalling targets responding to light are associated with changes to transcription^1^. Indeed, during the evolution of C_4_ photosynthesis, genes important for the C_4_ cycle appear to have acquired mutations that allowed integration into the light responsive transcriptional networks that operate in C_3_ species^2,3^. However, in C_3_ Arabidopsis during de-etiolation protein synthesis (translation) is also reconfigured in response to light^4,5,6^. Although, in C_4_ maize translational regulation has been associated with the cell preferential accumulation of enzymes required for C_4_ photosynthesis^7,8^ to our knowledge there have been no detailed studies on translational efficiency during photomorphogenesis in C_4_ species. In particular, we sought to establish whether control of genes used specifically in the C_4_ pathway is modified such that their translation becomes fully co-ordinated with canonical photosynthesis genes. To better understand how translation is reprogrammed during de-etiolation and whether this is altered during the evolution of C_4_ photosynthesis we undertook paired RNA-Seq and Ribosome Profiling (Ribo-Seq) on C_3_ rice and C_4_ sorghum to define and decouple light-dependent translational responses and RNA turnover (**Figure 1**).

**Figure 1:**
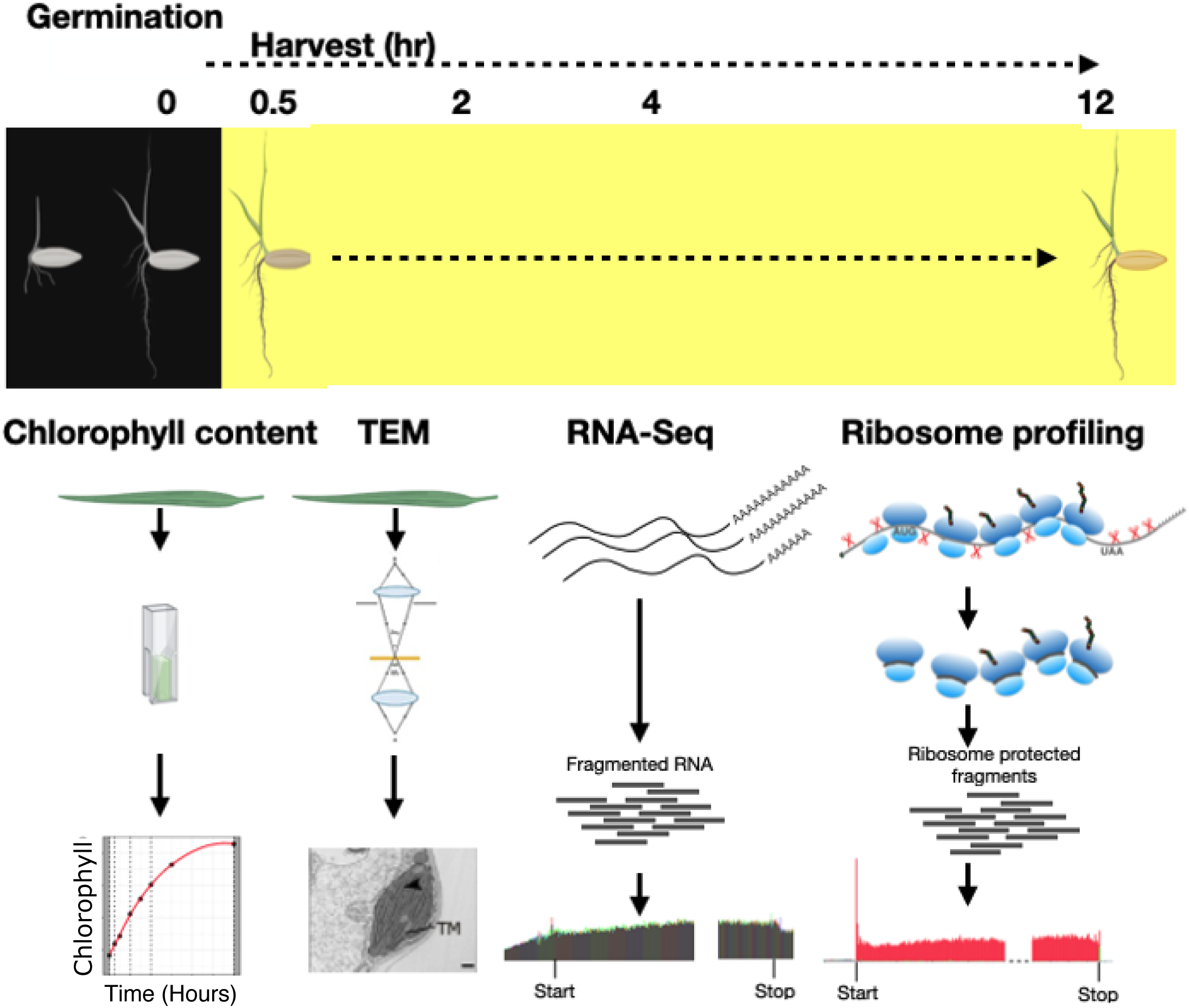
Schematic of experimental pipeline. Dark-grown seedlings of C_3_ *O. sativa* and C_4_ *S. bicolor* were exposed to light and the dynamics of chlorophyll accumulation as well as chloroplast ultrastructure by Transmission Electron Microscopy (TEM) defined. To understand the importance of changes in translation for these processes both RNAseq and ribosome profiling were undertaken.

## Results

### De-etiolation activates transcriptional and translational machinery in C_3_ rice

Prior to light exposure highly structured prolamellar bodies were evident in etioplasts of C_3_ rice (**Figure 2A, Supplemental Figure 1**). After 0.5 h of light exposure these had started to dissipate and by subsequent timepoints, development of thylakoid membranes with granal stacks was evident. Starch granules indicative of net photosynthesis were visible by 12 h. Concomitant with this elaboration of chloroplast structure, chlorophyll content accumulated in a logistic manner immediately after light exposure (**Figure 2B**). Biogenesis of chloroplasts was underpinned by accumulation of transcripts encoding proteins required for C_3_ photosynthesis (**Figure 2C**). These included transcripts encoding light harvesting complexes and the Calvin-Benson-Bassham cycle, but as would be expected not those derived from orthologs of C_4_ photosynthesis genes (**Figure 2C, Supplemental Figure 2**). Our results therefore suggest 12 hours of light was sufficient to capture events associated with chloroplast biogenesis and the dynamics of photosynthesis gene expression underlying this response.

**Figure 2:**
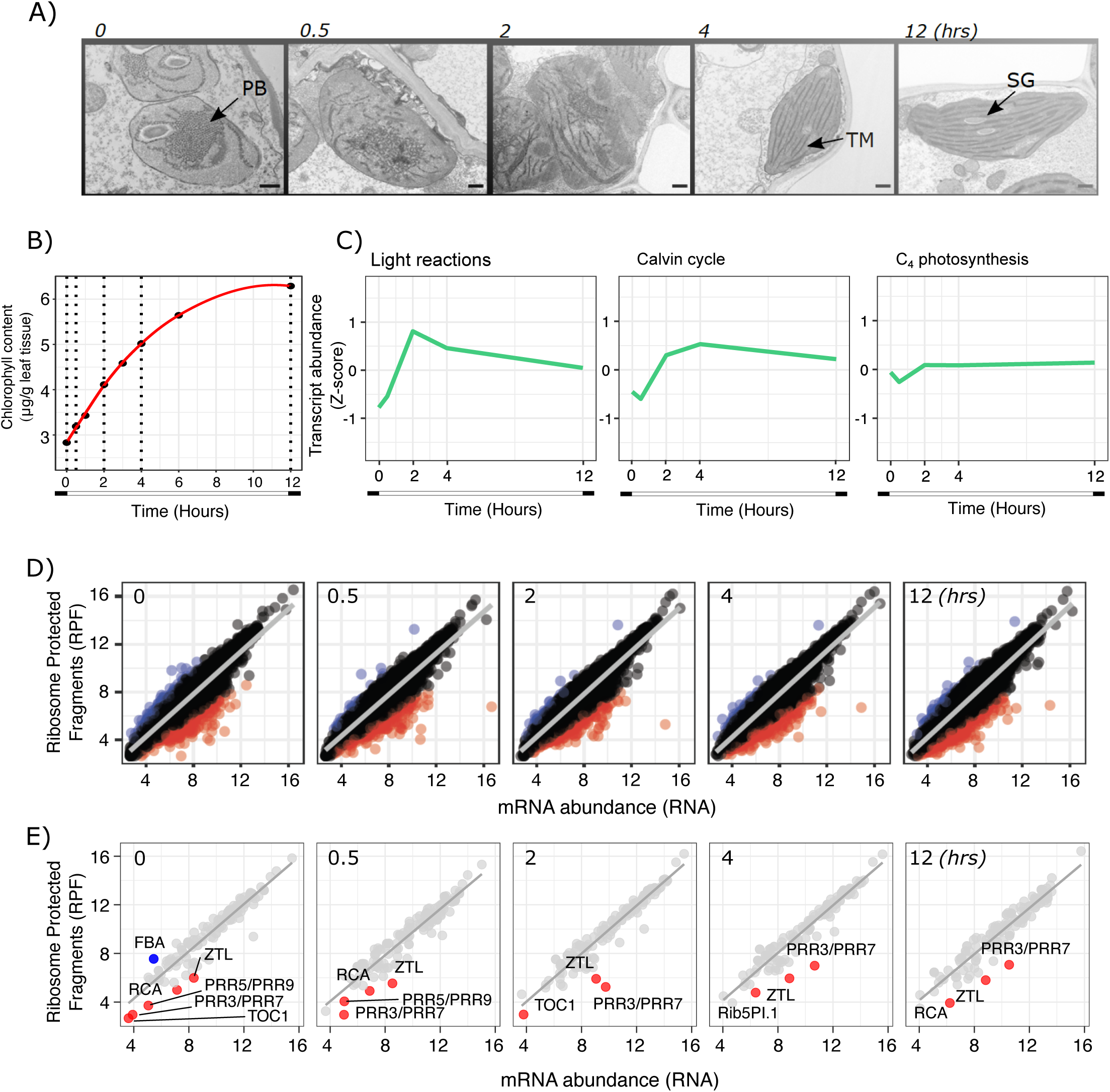
Dynamics of de-etiolation in rice (A) Transmission electron micrographs at 0, 0.5, 2, 4 and 12 hrs after light exposure, showing etioplasts to chloroplast development and assembly of the photosynthetic apparatus in rice mesophyll cells. Abbreviations: PB; Prolamellar body, TM; Thylakoid Membrane, SG; Starch Granule. Scale bars represent 500 nm. (B) Accumulation of chlorophyll after exposure to light. (C) Line plots depicting average changes in abundance of transcripts (z-scores) derived from genes associated with the light dependent reactions of photosynthesis, the Calvin-Benson-Bassham cycle and C_4_ photosynthesis. D) Variance Stabilized Counts (VST) for RNAseq (RNA) versus Ribo-Seq (RPF) after 0, 0.5, 2, 4 and 12 hrs of light exposure. A linear model was fitted to the data (p < 0.001). Blue and red dots represent genes not fitting a linear relationship (standard residuals > 2 (blue) or < –2 (red)). E) VST counts for RNA versus RPF from selected photosynthesis and light signalling genes. Blue and red dots represent genes not fitting a linear relationship (standard residuals > 2 (blue) or < –2 (red)). Abbreviations: FBS; Aldolase1, RCA; Rubisco activase, ZTL; Zeitlupe, PRR3,5,7; Pseudo-response-regulator, TOC1; Timing of cab expression 1.

Although the induction of light-mediated transcriptional networks is fundamental for establishment of the photosynthetic machinery^9^, the extent to which this is co-ordinated with protein synthesis is not fully defined. Ribo-Seq is a high-throughput technology that captures positions of ribosomes on mRNAs and so provides a highly-sensitive ‘snap-shot’ of global translational activity at any time point^10^ (**Figure 1**). When undertaken in parallel with RNA-Seq it provides insight into the extent to which increases in transcript abundance are coupled with protein synthesis. Quantification of the total number of ribosome protected fragments (RPF) provides a direct measurement of total protein synthesis, and the number of RPF per RNA molecule an estimate of translation efficiency (TE). In contrast with the read-size distribution from the RNA-Seq data that was indistinct, devoid of phasing and included reads from both 5’ and 3’ UTRs (**Supplemental Figure 3B**) Ribo-Seq generated profiles enriched in size classes of between 27 and 29 nucleotides, distinct phasing, and RPF were not detected in untranslated regions (UTRs, **Supplemental Figure 3A&B**). As has previously been reported in mouse*, Chlamydomonas reinhardtii* and *Arabidopsis thaliana* we also observed a larger footprint at the stop codon (**Supplemental Figure 3C)**. The latter characteristic has previously been associated with release factor rather than tRNA incorporation during translation termination^11,12^. It was also notable that as with data from *C. reinhardtii* and *A. thaliana*^12,13^ an enrichment of reads corresponding to start and stop codons was observed in the meta-transcriptomes due to preferential ligation of adaptors to 5’ Adenosine and 5’ Uridine start and stop codons respectively during library preparation (**Supplemental Figure 3B)**. We also detected the non-AUG) initiated upstream open reading frame for the *GDP-L-GALACTOSE PHOSPHORYLASE* (*GGP*) gene involved in L-ascorbate biosynthesis (**Supplemental Figure 4).** Non-canonical regulation of *GGP* is found in *Chlamydomonas reinhartii* as well as *Arabidopsis thaliana* and *Triticum aestivum*^14^. In summary, characteristics of the data from Ribo-Seq conformed to expectations and so with the RNA-Seq collected in parallel should allow robust analysis of any uncoupling between RNA levels and translation during de-etiolation.

We next analysed global translation profiles to determine the extent to which translational regulation during de-etiolation modified the transcriptional response. Principal Component Analysis performed on normalized counts from RNA-Seq (RNA) and Ribo-Seq showed that 39% of variance was explained by time exposed to light. The second principal component related to mRNA abundance and protein synthesis (**Supplemental Figure 5**). Correlation analysis between mRNA abundance and protein synthesis showed that they were strongly correlated indicating a tight coupling between mRNA abundance and translation (p < 0.001, **Figure 2D, Supplemental Figure 6**). However, residuals analysis revealed that the relationship between mRNA levels and protein synthesis (total RPF) was not linear for all translated genes (**Figure 2D&E**, **Supplemental Table 1**). For example, in the dark, one gene encoding fructose-1,6-bisphosphate aldolase (FBA, *LOC_Os10G08022*) which is important for gluconeogenesis during early seedling growth showed higher rates of protein synthesis than expected (**Figure 2E**). Gene enrichment analysis for genes showing a weaker coupling between protein synthesis and mRNA abundance in the dark (0 hrs) showed that these genes tended to be classified into categories related to organelle organization including “nucleus” and “membrane bounded organelles” (FDR < 0.05) (**Supplemental Figure 7**). In contrast, by 4 and 12 h after light exposure genes annotated as “cytoplasmic” tended to be translated more than would be predicted from their transcript abundance (**Supplemental Figure 7, Supplemental table 5**). It was noticeable that over the time course genes related to light signalling including the blue light photoreceptor Zeitlupe (ZTL) and also several circadian rhythm related genes in the PRR family (PRR3-7, PRR5-9 and TOC1) displayed lower rates of protein synthesis than was expected from mRNA abundance (**Figure 2E**).

### Rice photosynthesis transcripts with low abundance in the dark are efficiently translated

The above analysis suggested that in rice the translational programme operating in etiolated plants is disrupted by 30 minutes of light. To further explore these changes we compared Translational Efficiency (TE) with mRNA abundance. TE is defined as the ratio of counts derived from Ribo-Seq (RPF, i.e. protein synthesis) compared with those coming from total mRNA, and is therefore a measure of the proclivity of an mRNA molecule to interact with one or more ribosomes simultaneously. TE thus provides insight into the economics of translation (**Figure 3A**). Exposure to light for 30 mins had relatively little impact on coupling between TE and mRNA abundance with ninety-one percent of genes showing less than two-fold change compared to the dark period for both TE and mRNA abundance (**Figure 3B**). Of the remaining 9% of genes that did not correlate strongly, four percent showed two-fold higher TE in the dark but this was no longer detected 30 minutes after light (**Figure 3B, Supplemental table 4**). A similar trend was found for genes with higher TE after 0.5 hrs of light compared with the dark (**Supplemental Figure 8A**). Analysis of efficiently translated genes in the dark showed an enrichment in categories related to DNA packaging such as core histones and chloroplast proteins. This is consistent with the regression analysis for mRNA and total RPF as the residual levels of transcripts classified by the “chloroplastic” gene ontology term were higher in the dark compared with 0.5 h of light (**Figure 3C, Supplemental Figure 8B**). Among genes associated with the chloroplast we found transcripts encoding a signal recognition particle *OscpSRP43* (LOC_Os03g03990) which is essential for chloroplast development and photosynthesis^15^. Other genes included those encoding a Photosystem II core complex protein *psbY* (LOC_Os08g02630), the plastid specific ribosomal protein 6 (LOC_Os11g34940) and the Calvin-Benson-Bassham cycle protein *CP12* (LOC_Os01g19740) (**Supplemental Table 5)**. For genes that were less translated in the dark but rapidly acquired more ribosomes in response to light the only enriched category was defined by intracellular anatomical structures. For instance, the transcription factor *DOF12* (*LOC_Os03g07360*) known to be involved in defining leaf architecture (Wu *et al*., 2015) and perhaps associated with the morphological changes taking place during de-etiolation (**Supplemental Figure 8C, Supplemental table 7**). Overall, our results indicate the translational program of skotomorphogenic rice leaves is characterised by high translational efficiency for genes involved in chloroplast biogenesis. This is then temporally disrupted by light, before a tight coupling between TE and mRNA abundance is established.

**Figure 3:**
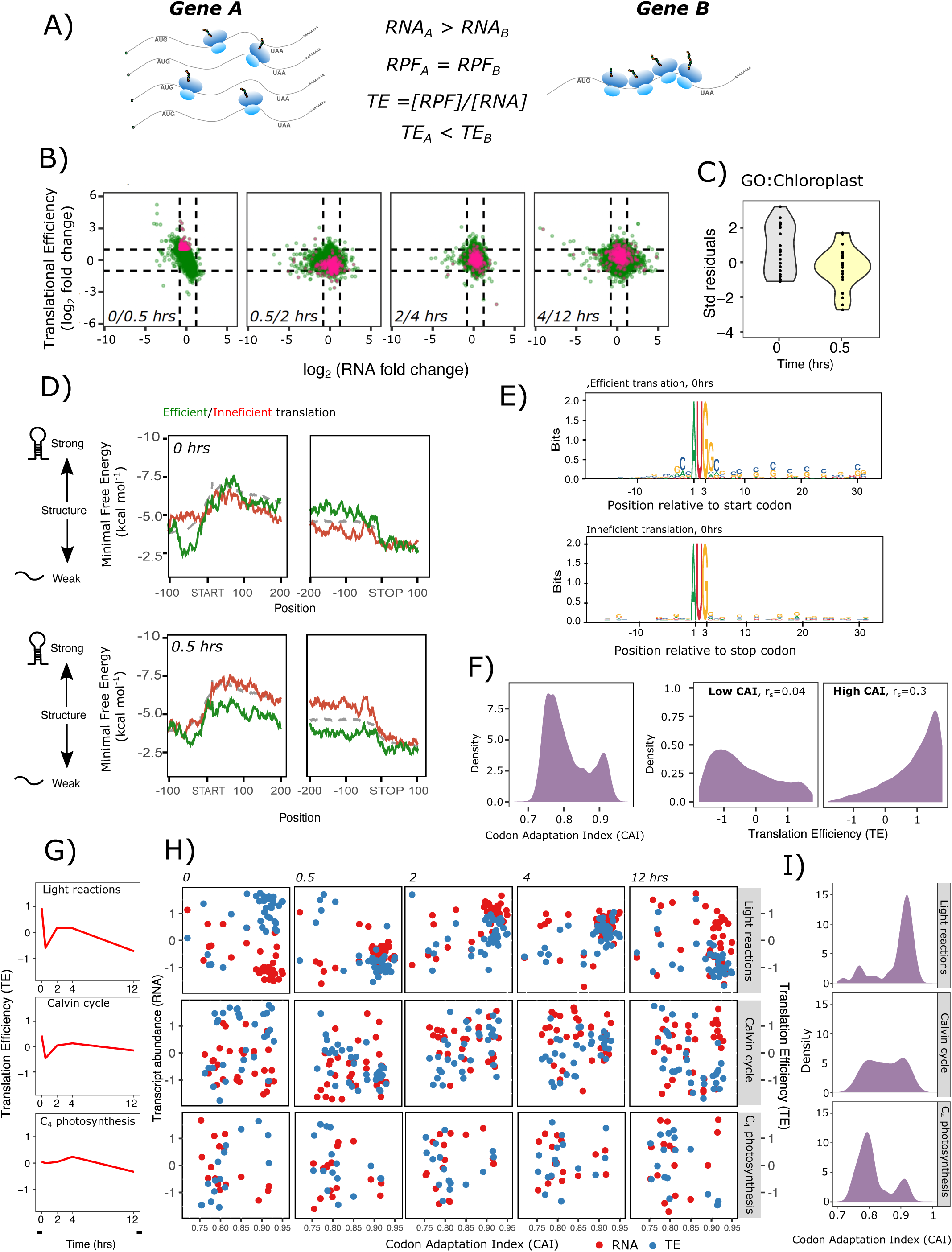
Translational dynamics during de-etiolation in rice. A) Schematic of how translation efficiency (TE) was defined. Gene B has a higher TE than Gene A because number of ribosome protected fragments per mRNA strand is higher. B) Log2 Fold changes for TE compared with total RNA levels. Pink dots reflect genes showing at least 2-fold increase in TE and no change in mRNA levels 30 mins after exposure to light and their position in the following timepoints. C) Violin plots representing the standard residuals at 0 and 0.5 h after illumination obtained from linear regression for genes whose Gene Ontology was annotated “Chloroplastic”. D) Predicted sliding-window minimal free energy of RNA secondary structures for nucleotides –100 to +200 from the start codon (left) and – 200 to +100 from stop codon (right). Grey dashed line represents mean of all expressed genes. Mean values for 1% of genes with the highest (efficient) and lowest (inefficient) TE shown in green and red respectively. E) Sequence logos for 1% of genes with the highest (efficient) and lowest (inefficient) TE at 0 h. F) Distribution of Codon Adaptation Index (CAI) across all transcripts analysed (left). Relationship between CAI and TE for genes in the dark for genes with CAI < 0.85 (center) or ≥ 0.85 (right). Pearson correlation between CAI and TE shown. G) Average translation efficiency (TE) shown as Z-scores for genes associated with the light dependent reactions of photosynthesis, the Calvin-Benson-Bassham cycle and C_4_ photosynthesis. H) mRNA abundance (red dots) and TE (blue dots) during de-etiolation. I) Distribution of CAI associated with the light dependent reactions of photosynthesis, the Calvin-Benson-Bassham cycle and C_4_ photosynthesis.

To better understand the translational response associated with photomorphogenesis we examined the following features for differentially translated transcripts: (1) RNA structure, which can interfere with all stages of translation; (2) initiation context, as initiation is a rate-limiting stage of translation; and (3) codon usage, which can impact efficiency of translational elongation. Plotting the local minimal free energy of RNA secondary structure along a 30-nucleotide sliding window revealed that the most efficiently translated mRNAs in the dark typically possessed unstructured 5’ UTRs **(Figure 3D and Supplemental Figures 9)**. However, after 0.5 h of light efficient translation was associated with a different class of mRNAs that possessed less structured coding sequence. These findings are consistent with two modes of translational regulation allowing control over ribosomal scanning/initiation and elongation in dark and light respectively **(Figure 3D, Supplemental Figure 9)**. As initiation is widely considered a rate-determining step for efficient translation and we also observed preference for unstructured 5’UTRs in efficiently translated mRNAs in the dark, we also investigated sequence identity surrounding the start codon (i.e. Kozak consensus) to determine whether efficiently translated mRNA are subject to efficient 5’UTR scanning as well as initiation. However, while we revealed a rice-specific Kozak consensus, where instead of A/GNN_AUG_G, efficiently translated rice mRNAs were generally enriched in guanidine at +6 and cytosine at the –2 and +5 positions (**Figure 3E, Supplemental Figure 10**), we did not observe significant differences in initiation context associated with transcripts showing altered TE in response to light.

Preferential translation of mRNAs with less structured coding sequence after thirty minutes of light could be associated with translational elongation through a reduced need to unwind RNA. We investigated whether this effect is also compounded with efficient codon usage. Codon adaptation index (CAI) is a measure of codon bias and when this was plotted for all translated transcripts a bimodal distribution with peaks around 0.76 and 0.92 was apparent **(Figure 3F)**. Further investigation of the relationship between TE and CAI during light transition revealed that efficiently translated transcripts in the dark were generally associated with high CAI **(Figure 3F)**. However, this correlation was lost upon light induction **(Supplemental Figure 11)**. From these data, we propose that within first 30 minutes of light induction TE is mainly controlled during elongation.

We next focused on transcripts associated with photosynthesis as we observed high TE and low transcript abundance of genes associated with light-dependent reactions in the dark (**Figure 2C and 3G**). For this gene set, TE decreased after thirty minutes of light because transcript abundance increased, before rising again at 2 and 4 hours of light. This behaviour was also observed for genes encoding proteins of the Calvin-Benson-Bassham cycle (**Figure 3G**). In contrast, TE of genes encoding proteins associated with C_4_ photosynthesis were not affected by light (**Figure 3G**). Further investigation revealed genes involved in the light-dependent reactions of photosynthesis with a high CAI (>0.85) typically had low transcript abundance but high TE in the dark **(Figure 3H-I)**. After 30 minutes and two hours of light, transcript derived from these genes with high CAI had increased whilst TE had dropped **(Figure 3H)**. There was little change after 4 hrs and by 12 hrs of light in transcript abundance and TE of these genes had dropped at 12hr **(Figure 3H)**. These co-ordinated changes to transcript abundance and TE for genes with high CAI were not as evident for components of the Calvin-Benson-Bassham or C_4_ cycles (**Figure 3H**). In fact, when the relationship between CAI and these gene categories was plotted there was a clear enrichment of high CAI for genes associated with the light dependent reactions of photosynthesis, a more balanced distribution for genes of the Calvin-Benson-Bassham cycle, and a higher proportion of C_4_ related genes with low CAI **(Figure 3I)**. These data indicate that in rice most of the genes associated with light dependent reactions of photosynthesis and a reduced proportion of Calvin-Benson-Bassham cycle appear to have evolved an optimised codon usage, and despite low transcript abundance are efficiently translated in the dark. These characteristics are not apparent for rice genes whose orthologs are co-opted into the C_4_ cycle in C_4_ plants.

We conclude that TE dynamics during photomorphogenesis of rice is determined by at least three processes. First, efficiently transcripts in the dark tend to possess 5’UTRs with relaxed secondary structure, indicative of efficient 43S ribosomal scanning prior to initiation^16^. Second, upon transfer to light, coding region for transcripts with high TE typically have a relaxed structure which facilitates efficient translation elongation by circumventing need for additional helicase activities during elongation. Lastly, genes encoding components of the light dependent reactions of photosynthesis have evolved to possess optimised codon usage for efficient translational decoding in the dark.

### C_4_ cycle genes in *Sorghum bicolor* are efficiently translated in the dark

We next investigated whether genes encoding proteins of the C_4_ pathway in sorghum have co-opted this translational control of photosynthesis genes that operates during photomorphogenesis of C_3_ rice. Over the same de-etiolation time-course as used above, chlorophyll accumulated and chloroplast biogenesis was induced in sorghum **(Figure 4A and B, Supplemental Figure 12**). Dimorphic chloroplasts in mesophyll and bundle sheath cells^17^ were observed by 4 hrs of light with granal stacks preferentially observed in the mesophyll and starch granules most evident in the bundle sheath (**Figure 4B**). This assembly of the photosynthetic apparatus was accompanied by accumulation of transcripts encoding components of the light dependent reactions of photosynthesis, Calvin-Benson-Bassham cycle but also genes required for the C_4_ pathway (**Figure 4C, Supplemental Figure 13**). There was a strong linear relationship between mRNA abundance and protein synthesis activity (RPF) (**Supplemental Figure 14, 15 & 16A)**. As with rice, time after exposure to light was the main component for driving variance (42%) (**Supplemental Figure 15**), and light-signalling and circadian clock genes such as *ZTL*, *PRR3-7, TOC1, CCA/LHY* had lower rates of protein synthesis than would be expected from the linear model predicted from all RNA abundance and RPF data (**Supplemental Figure 16B**). Moreover, 7.5% of translated genes showed a two-fold change in TE without a significant change in transcript abundance and this was most noticeable during the transition from dark to light (**Supplemental Figure 16**). Gene Ontology analysis for genes with higher TE in the dark than 30 minutes of light showed an enrichment in chloroplastic genes (**Supplemental Figure 17B & 18**). However, the identity of chloroplastic genes enriched in sorghum were different to those found in rice with for example genes encoding proteins related with NAD-binding and localized in the stroma as well as ribosomes being enriched (**Supplemental Table 10**). For genes with higher TE at 0.5 hrs of light compared with the dark we observed a similar trend to rice where this difference was lost by later timepoints (**Supplemental Figure 17, Supplemental Table 11**).

**Figure 4:**
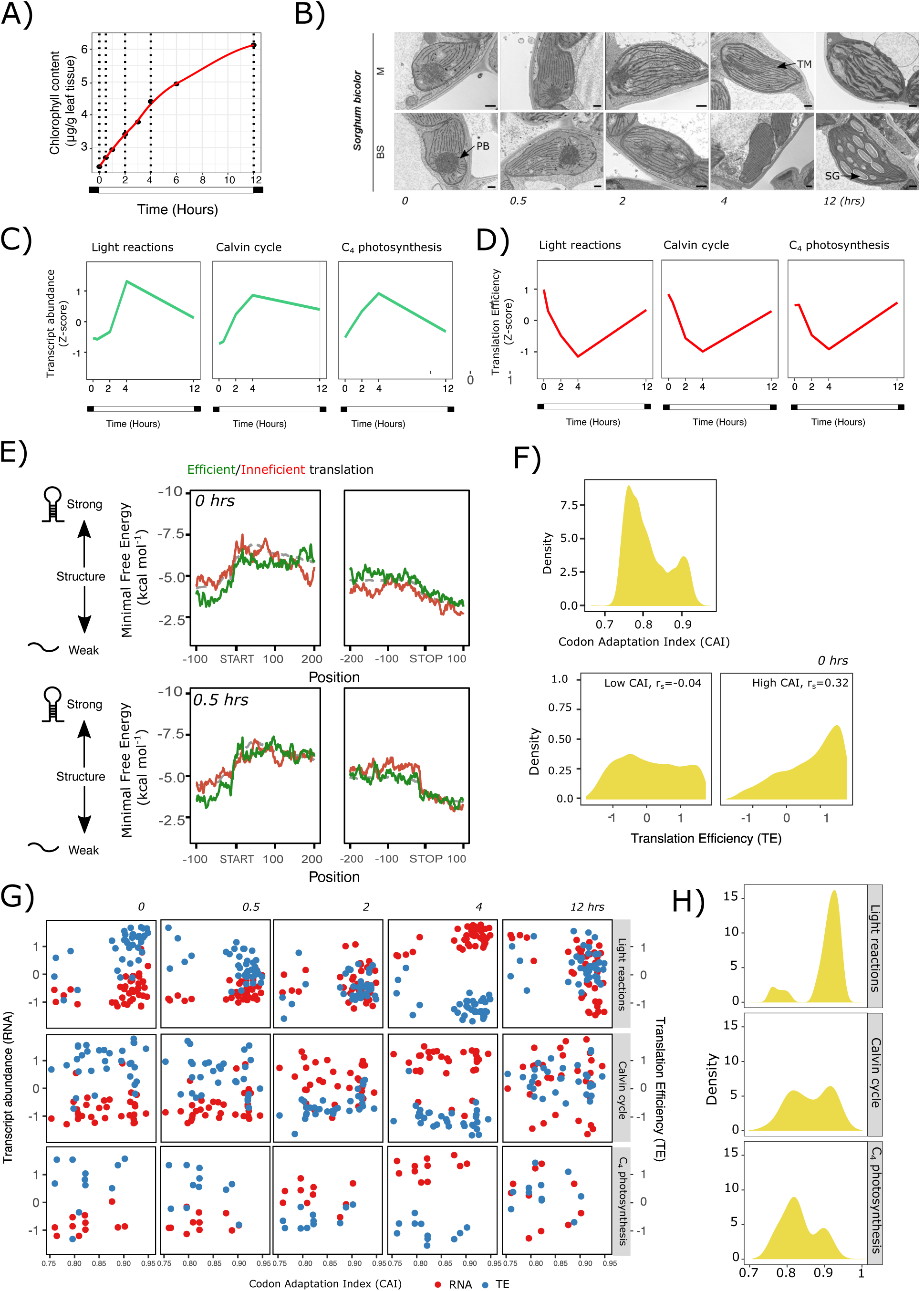
Dynamics of de-etiolation in sorghum (A) Accumulation of chlorophyll after exposure to light. (B) Transmission electron micrographs at 0, 0.5, 2, 4 and 12 hrs after light exposure, showing etioplast to chloroplast development and assembly of the photosynthetic apparatus in sorghum mesophyll (M) and bundle sheath (BS) cells. PB; Prolamellar bodies, TM; Thylakoid Membrane, SG; Starch granule. Scale bars represent 500 nm. (C) Line plots depicting average changes in abundance of transcripts (z-scores) derived from genes associated with the light dependent reactions of photosynthesis, the Calvin-Benson-Bassham cycle and C_4_ photosynthesis. D) Average translation efficiency shown as Z-scores for genes associated with the light dependent reaction of photosynthesis, the Calvin-Benson-Bassham cycle and C_4_ photosynthesis. E) Predicted sliding-window minimal free energy of RNA secondary structures for nucleotides –100 to +200 from the start codon (left) and – 200 to +100 from stop codon (right). Grey dashed line represents mean of all expressed genes. Mean values for 1% of genes with the highest (efficient) and lowest (inefficient) TE shown in green and red respectively. F) Distribution of Codon Adaptation Index (CAI) across all transcripts analysed (top). Relationship between CAI and TE for genes in the dark for genes with CAI < 0.85 (bottom left) or ≥ 0.85 (bottom right). Pearson correlation between CAI and TE shown. G) mRNA abundance (red dots) and TE (blue dots) during de-etiolation. H) Distribution of CAI for genes associated with the light dependent reactions of photosynthesis, the Calvin-Benson-Bassham cycle and C_4_ photosynthesis.

**Figure 5:**
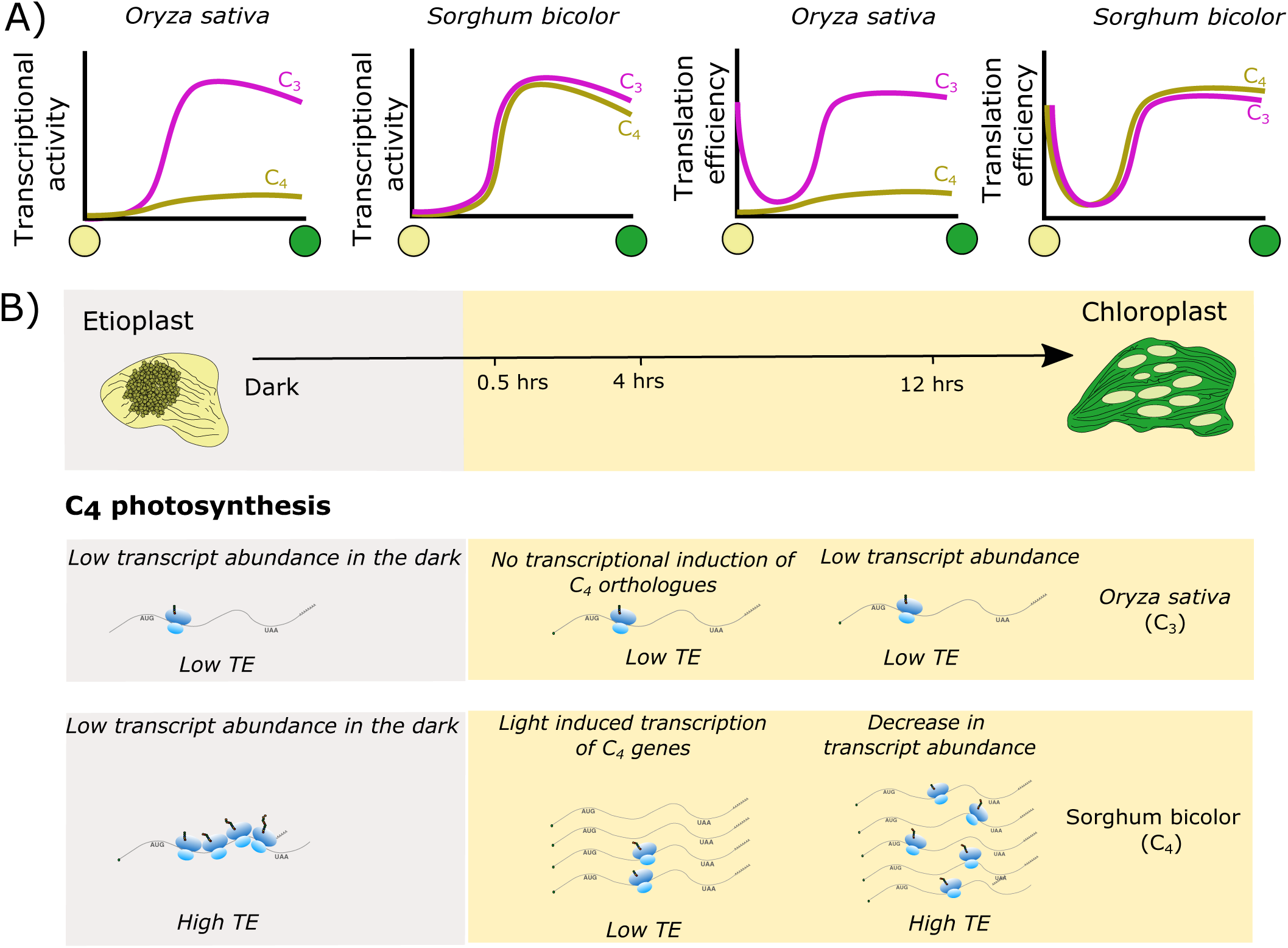
Model for transcriptional and translational dynamics of photosynthesis genes during de-etiolation. (A) In the dark, low transcription rates for genes of the Calvin-Benson-Bassham cycle and light dependent reactions of photosynthesis are compensated with relatively high translation efficiencies (TE) in both C_3_ rice and C_4_ sorghum. Light triggers activation of the transcriptional machinery for these genes reducing the number of ribosomes engaged in translation per mRNA molecule. Towards the end of de-etiolation protein synthesis activity is coupled with transcript abundance resulting in an increase in TE (B) In rice, these patterns were not observed for C_4_ genes. However, in sorghum C_4_ genes follow the same pattern as other photosynthesis genes implying they have been re-wired to pre-existing transcriptional and translational networks.

Analysis of the dynamics of TE for photosynthetic genes revealed that in addition to genes encoding components of the light dependent reactions of photosynthesis showing a rapid reduction in TE after exposure to light, this was also the case for those associated with the Calvin-Benson-Bassham cycle and C_4_ photosynthesis (**Figure 4D**). Like rice, there was no clear differences in Kozak initiation consensus sequence between differentially translated transcripts in either dark or light (**Supplemental Figure 20**). In contrast with rice, there was no compelling evidence for transcripts with high TE in the dark having lower structure of either UTRs or coding regions suggesting that in sorghum additional mechanisms such as recruitment of *trans-*factors facilitate translational regulation in the dark (**Figure 4E, Supplemental Figure 19**). However, efficiently translated sorghum transcripts in the dark tended to possess high codon usage (CAI) (**Figure 4F and Supplemental Figure 21**). As with rice, this was a global observation and was particularly evident for genes associated with the light dependent reactions of photosynthesis (**Figure 4F-H**).

Genes associated with the light dependent reactions of photosynthesis in sorghum with high CAI showed a gradual reduction in TE upon light exposure that was then reversed at 12 hrs of light exposure **(Figure 4G)**. In contrast, abundance of transcript derived from these genes with high CAI was low in the dark but rose steadily from 2 hrs onwards **(Figure 4G)**. Similar to rice, it was clear that the majority of sorghum genes associated with the light dependent reactions of photosynthesis had high CAI, while the proportion of genes with high CAI was lower for Calvin-Benson-Basham genes and even lower for C_4_ cycle genes **(Figure 4H)**. Among those genes we observed that in both sorghum and rice, a gene encoding the small sub-unit of RuBisCO (*SOBIC.005G042000* and *LOC_Os12G17600*) had high levels of CAI (**Supplemental Figure 22**). It is known that translation elongation plays an important role in *RbcS* regulation in other C_4_ species such as *Amaranthus hypochondriacus*^8,18,19^ suggesting codon adaptation could be contributing to this type of regulation in C_3_ and C_4_ species.

While there was no clear correlation between CAI and TE for C_4_ genes the behaviour of C_4_ cycle genes in sorghum paralleled the translational behaviour of light responsive genes **(Figure 4G)**. Among the C_4_ genes with high CAI values we found *CA* and *PEPC.* These genes have also high CAI score in rice, and whilst both genes were efficiently translated in dark in sorghum this was not the case in rice (CAI*^CA.sorghum^*= 0.88, CAI*^PEPC.sorghum^* =0.90 and CAI*^CA.rice^*= 0.915, CAI*^PEPC.rice^* =0.917, **Supplemental Figure 22)** suggesting this feature is important but not sufficient for efficient translation of C_4_ cycle genes in the dark in sorghum.

## Discussion

Depending on the light environment seedlings adopt different developmental programmes. In the dark, a skotomorphogenic plant will remobilise reserves to allow hypocotyl elongation and root development to prepare for emergence through the soil and onset of light and autotrophy. When light is absorbed, a complex signal cascade mediated by photoreceptors triggers photomorphogenesis through activation of transcriptional networks^1,20^. In the case of C_4_ species, there is an additional requirement that C_4_ genes are incorporated into these light responsive regulatory networks. It is appears that during evolution this has occurred at least in part through the acquisition of light responsive *cis*-elements^3,21^. Although photomorphogenesis in both C_3_ and C_4_ leaves is highly dependent on remodelling of transcriptional activities, it is clear that post-transcriptional regulation is also crucial role to allow efficient assembly of the photosynthetic machinery^22^. To explore the dynamics of translation during the establishment of C_3_ and C_4_ photosynthesis we applied Ribo-Seq and RNA-Seq during de-etiolation of *O. sativa* and *S. bicolor*. Consistent with previous observations of the C_3_ plant *Arabidopsis thaliana*^5^ mRNA turnover seems to be the main determinant in the regulation of most photosynthesis related genes upon light induction. We conclude that this feature appears conserved between C_3_ and C_4_ species.

However, our analysis also shows that for many light responsive genes the relationship between translational responses and mRNA turnover was not linear. The relatively low rates of protein synthesis for a group of clock regulators (ZTL, PPRs and TOC1) could allow different elements of the clock to be fully integrated with activation of the photosynthetic machinery. In both species after 12 hours of light a steady state relationship between RNA abundance and translation had been re-established. Skotomorphogenesis operates with limited resources and so once light is available, photosynthetic competence must be achieved quickly. Consistent with this, non-photosynthetic etioplasts contain protein complexes required for electron transport and protein synthesis^23^ implying some basal activity of photosynthesis related genes that is independent of light signalling networks takes place in the dark. Our data show that in the dark photosynthesis-related genes are efficiently translated despite low transcript abundance, and so indicate a compensatory mechanism maximising production of protein from these low levels of transcripts. As the light environment becomes favourable for activation of light-signalling cascades, photosynthesis genes become more dependent on transcriptional activity. Interestingly, the identity of genes most influenced by these changes as well as the duration of this translational compensation differed between rice (C_3_) and sorghum (C_4_) possibly as a consequence of different protein stoichiometries needed for the functioning of the photosynthetic apparatus. Ribosome profiling has been undertaken at multiple developmental stages^24^ and between mesophyll and bundle sheath cells of C_4_ maize^7,24^. The main focus of these analysis was on chloroplast genes, and it was concluded that changes in mRNA abundance were the major control of protein production, with changes in translational efficiency allowing fine-tuning^24^. Moreover, It was also shown that translation efficiency for genes encoding PSII core subunits such as psbA, psbB, psbC and psbD had a significant impact on preferential gene expression in the mesophyll^24^. Subsequent analysis of maize chloroplasts from green seedlings experiencing light-to-dark and dark-to-light shifts showed enhanced ribosome occupancy for *psbA* in response to light. While other chloroplast mRNAs showed similar ribosome occupancy during these transitions, translation efficiency was higher due to an increase in elongation rate^25^. Together with our findings, these results suggest that in C_3_ and C_4_ cereals genes involved in the light-dependent reactions of photosynthesis (either chloroplastic or nuclear encoded) are tightly controlled at the translational level. This is apparent despite the fact chloroplast translation is compartmentalised and performed by prokaryotic-type ribosomes, and nuclear genes are translated by cytosolic eukaryotic type ribosomes.

During photomorphogenesis while photosynthesis genes from both rice and sorghum are regulated at the level of translation, it is striking that the underpinning molecular mechanisms varied between the species. It is possible that these differences are caused by divergence between the lineages since their last common ancestor. Notably, we observed efficiently translated rice transcripts in the dark tended to possess unstructured 5’ UTRs, and although not the case in sorghum, transcripts in rice with less structured coding regions were preferentially translated after exposure to light. It is possible that in rice in the dark transcripts that enable efficient 43S ribosomal scanning process are preferred whilst transcripts with unstructured coding regions are preferentially translated by the 80S ribosome upon light induction^16,26,27^. We propose that processes independent of RNA structure operate in sorghum during the dark to light transition, potentially through recruitment of additional *trans*-factors, either to the mRNA or the 43S/80S ribosome to facilitate efficient 5’ UTR scanning and/or translational elongation.

In both rice and sorghum it appears that codon optimality plays a major role in efficient translation of photosynthesis genes, especially those involved in the light-dependent reactions when plants are skotomorphogenic. Upon light perception, transcriptional induction is initiated and transiently exceeds the rate of translation before coupling is restored. As this was observed in rice and sorghum we propose that this is a conserved mechanism to optimise assembly of the photosynthetic apparatus once light is perceived. Post transcriptional control of C_4_ gene expression has been established in the past mainly in the context of cell-preferential gene expression^7,8,18,19,24,28–30^ and chloroplast differentiation^24^. In maize at least 31 genes associated with the C_4_ cycle showed differences in translation between mesophyll and bundle sheath cells^7^. Of these, *Ribose-5-phosphate isomerase* (R5PI) was found to be subject to translational control in maize but also the independent C_4_ lineage *Setaria viridis*^31^. Our work shows R5PI orthologs for both sorghum and rice had high translation efficiency in the dark that was associated with high codon use efficiency. Together, these findings imply that genes important for photosynthesis are regulated at the level of translation during de-etiolation and seedling development but also in the mature C_4_ leaf. While differential use of AUG versus non-AUG initiation codons in upstream open reading frames (uORFs) has been linked to cell-type dependent translation^7^ we did not find compelling evidence that uORFs or non-AUG initiation influence translation of photosynthesis related genes in rice or sorghum. This included orthologs of GUANINE NUCLEOTIDE EXCHANGE FACTOR (GEF), ADP-GLUCOSE PYROPHOSPHORYLASE LARGE SUBUNIT 1 (AGPLL1), CBL-INTERACTING PROTEIN KINASE 3 (CIPK3), and CHLOROPLAST UNUSUAL POSITIONING 1 (CHUP1) for which this has been reported for maize^7^. Whilst, in rice and sorghum it is possible that greater sequencing depth would reveal such phenomena for these genes, and the coverage was too low to determine the precise start codon utilised, we did find evidence for non-AUG initiation of Calvin-Benson-Bassam cycle regulatory protein CP12 in sorghum (Supplementary Figure 23).

In summary, this analysis shows that during photomorphogenesis and assembly of the photosynthetic apparatus tight control of translation of photosynthesis genes is apparent in both rice and sorghum. This regulation of translation has become more widespread in sorghum to include genes encoding proteins of the C_4_ cycle such that translation and transcription of canonical photosynthesis and also C_4_ pathway genes are co-ordinated. We conclude that during the evolution of the C_4_ pathway, genes encoding components of the C_4_ cycle have acquired mechanisms controlling translation of photosynthesis genes during photomorphogenesis in the ancestral C_3_ state.

## Materials and methods

### Growth of plant material, chlorophyll analysis and microscopy

Sterile rice (*Oryza sativa*) and sorghum (*Sorghum bicolor*) seeds were germinated on moist filter paper in the dark at 30°C for 24h. Germinated seeds were then transferred to 1:1 mixture of top-soil and sand for five days in a controlled environment growth room set at 28°C during the day and 25°C at night, with a relative humidity of 60%. Photomorphogenesis was induced by exposure to 350 μmol m^-2^s^-1^ white light photon flux density and photoperiod set to 12 hrs. Whole seedlings were harvested at 0.5, 2, 4 and 12 hrs after illumination (starting at 6:00am with the light cycle set from 6:00 to 18:00). Tissue was flash-frozen in liquid nitrogen and stored at –80°C before processing for RNA-SeqRNA-Seq and Ribo-Seq analysis.

For analysis of total chlorophyll content de-etiolating rice seedlings were flash-frozen at 0, 0.5, 2, 4 and 12 hrs after exposure to light. 100 mg of tissue was used for each extraction. Tissue was suspended in 1ml of 80% (v/v) acetone at 4°C for 10 minutes. The tube was then centrifuged at 13,250 g for 5 minutes and supernatant removed. The pellet was resuspended in 1 ml of 80% (v/v) acetone at kept at 4°C for 10 minutes prior to re-centrifugation. Supernatant from the two extractions were combined, and 1 ml transferred to a cuvette to allow absorbance to be measured in a spectrophotometer at 663.8 nm and 646.6 nm. The total chlorophyll content was analysed by using the equation derived by (Porra et al., 1989)^32^.

### Transmission Electron Microscopy

Leaf samples from dark-grown seedlings of *Oryza sativa* and *Sorghum bicolor* (youngest leaf at the three-leaf stage was chosen) were harvested for electron microscopy at 0, 0.5, 2, 4 and 12 hrs after light induction. Small leaf segments (∼2 mm^2^) were excised with a razor blade and immediately fixed in 2% (v/v) glutaraldehyde and 2% (w/v) formaldehyde in 0.05 – 0.1 M sodium cacodylate (NaCac) buffer (pH 7.4) containing 2 mM calcium chloride. Samples were vacuum infiltrated overnight, washed 5 times in 0.05 – 0.1 M NaCac buffer, and post-fixed in 1% (v/v) aqueous osmium tetroxide, 1.5% (w/v) potassium ferricyanide in 0.05 M NaCac buffer for 3 days at 4°C. After osmication, samples were washed 5 times in deionized water and post-fixed in 0.1% (w/v) thiocarbohydrazide for 20 min at room temperature in the dark. Samples were then washed 5 times in deionized water and osmicated for a second time for 1 h in 2% (v/v) aqueous osmium tetroxide at room temperature. Samples were washed 5 times in deionized water and subsequently stained in 2% (w/v) uranyl acetate in 0.05 M maleate buffer (pH 5.5) for 3 days at 4°C, and washed 5 times afterwards in deionized water. Samples were then dehydrated in an ethanol series, transferred to acetone, and then to acetonitrile. Leaf samples were embedded in Quetol 651 resin mix (TAAB Laboratories Equipment Ltd) and cured at 60°C for 2 days. Ultra-thin sections were cut with a diamond knife using a Leica Ultracut microtome and collected on copper grids and examined in a FEI Tecnai G2 transmission electron microscope (200 keV, 20 μm objective aperture). Images were obtained with an AMT CCD camera.

### Ribosome profiling and corresponding RNA-Seq

Samples were processed in similar manner as described by Chung et al 2017. Briefly, frozen leaf materials were pulverised in liquid nitrogen with pre-frozen buffer (20mM Tris-Cl pH 7.5, 140mM KCl, 5mM MgCl_2_, 0.5% NP40 (v/v), 1% Triton X-100 (v/v) and 5% (w/v) sucrose, 100µg/ml cyclohexamide and 100µg/ml chloramphenicol). The frozen powder was thawed and clarified by centrifugation at 4°C, 11200g for five minutes prior to dividing the lysate for either nuclease foot-printing with RNaseI for generation of the ribosome profiling libraries or total RNA extraction followed by removal of rRNA with RiboZero (plant leaf kit, Illumina) for the corresponding RNA-Seq library.

Libraries were sequenced with the NextSeq500 platform using 75 single read high-output v2 kit (TG-160-2005) and the reads were trimmed using the fastX toolbox and mapped with Bowtie 1 to the *Oryza sativa* and the *Sorghum bicolor* transcriptomes from Phytozome (versions 323.v7.0 and v3.1.1 respectively as of 2018). The mapped file was further processed with riboSeqR (Chung et al 2015) for *de novo* ORF detection. As our primary focus was on translated mRNAs and its regulation, we followed a filtering strategy by (a) selecting annotated open-reading frames that were supported by ribosome profiling data with in-frame AUG start and stop codons (UGA, UAA, UAG) where at least 50 in-frame reads mapping to at least ten unique locations across the mRNA (Chung et al 2015, 2017, 2020) and (b) parallel extraction of RNA-Seq reads as well as in-frame RPF reads within the defined ORF from (a). ORFs with overwhelmingly high transcript abundance but very little translation (i.e. TE >= 0.02) were also removed. This resulted in 9406 ORFs for *Oryza sativa* and 7858 ORFs for *Sorghum bicolor* for differential analysis with riboSeqR. The ORFs with TE < 0.02 (n= 41) were almost exclusively putative proteins associated with the photosynthesis apparatus or proteins with unknown function.

Minimal free energy (MFE) of RNA secondary structures was predicted by RNAFold from the ViennaRNA package (Lorenz et al. 2011). To relate predicted structure strength with position on the sequence, MFE for 30 nucleotide sliding window was calculated^33^. The mean MFE of a specified group of genes was plotted at each nucleotide position relative to the CDS start or end. Codon adaptive index (CAI) for all coding regions was calculated with EMBOSS^34^.

## Data availability

Raw and processed data are available from RiboSeq/RNASeq series (ArrayExpression accession E-MTAB-13489 and E-MTAB-13488).

## Code availability

Software package riboSeqR^61^ used in this study is opensource under BioConductor. Customised scripts used for this project are available upon request.

## Acknowledgements

This work was supported by a BBSRC DTP studentship to F.L., Medical Research Council Fellowship [MR/R021821/1], BBSRC project grants [BB/X001261/1, BB/V017780/1 and BB/V006096/1] to B.Y.W.C., and CEX2019-000902-S funded by MCIN/AEI/10.13039/501100011033 from the CERCA Programme/Generalitat de Catalunya to I.RL., as well as BBP0031171 and ERC Grant Revolution RG80867 to J.M.H. T.B.S. was supported by a Swiss National Science Foundation (SNSF) Early Postdoc Mobility Fellowship (P2EZP3_181620) and an EMBO Long-Term Fellowship (ALTF 531-2019). We thank Karin H. Müller and Filomena Gallo from the Cambridge Advanced Imaging Centre for the electron microscopy sample preparation as well as the support during the image acquisition. For the purpose of open access, the authors have applied a Creative Commons Attribution (CC BY) licence to any Author Accepted Manuscript version arising from this submission.

## Author contributions

I.RL., B.Y.W.C. and J.M.H. conceived the research and designed experiments. B.Y.W.C. and I.RL. generated Ribo-Seq and RNA-Seq libraries. T.B.S. performed TEM and P.S. performed chlorophyll quantification. I.RL. and F.L. performed bioinformatics analysis. I.RL., B.Y.W.C. and J.M.H. wrote the manuscript.

