## Supplementary Figures for "Pervasive translational control of photosynthesis genes during photomorphogenesis is acquired by C_4_ genes"

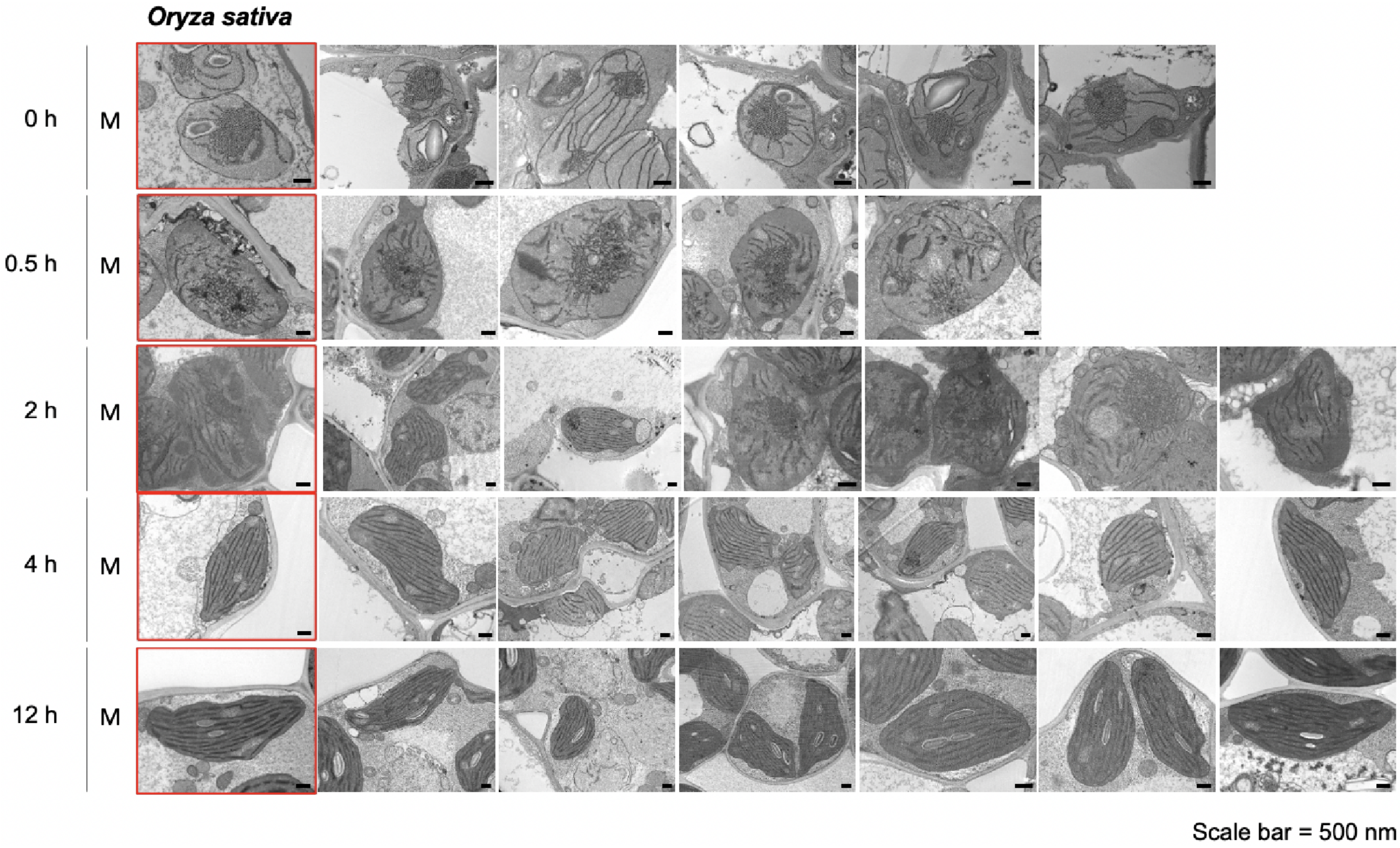


Figure S1

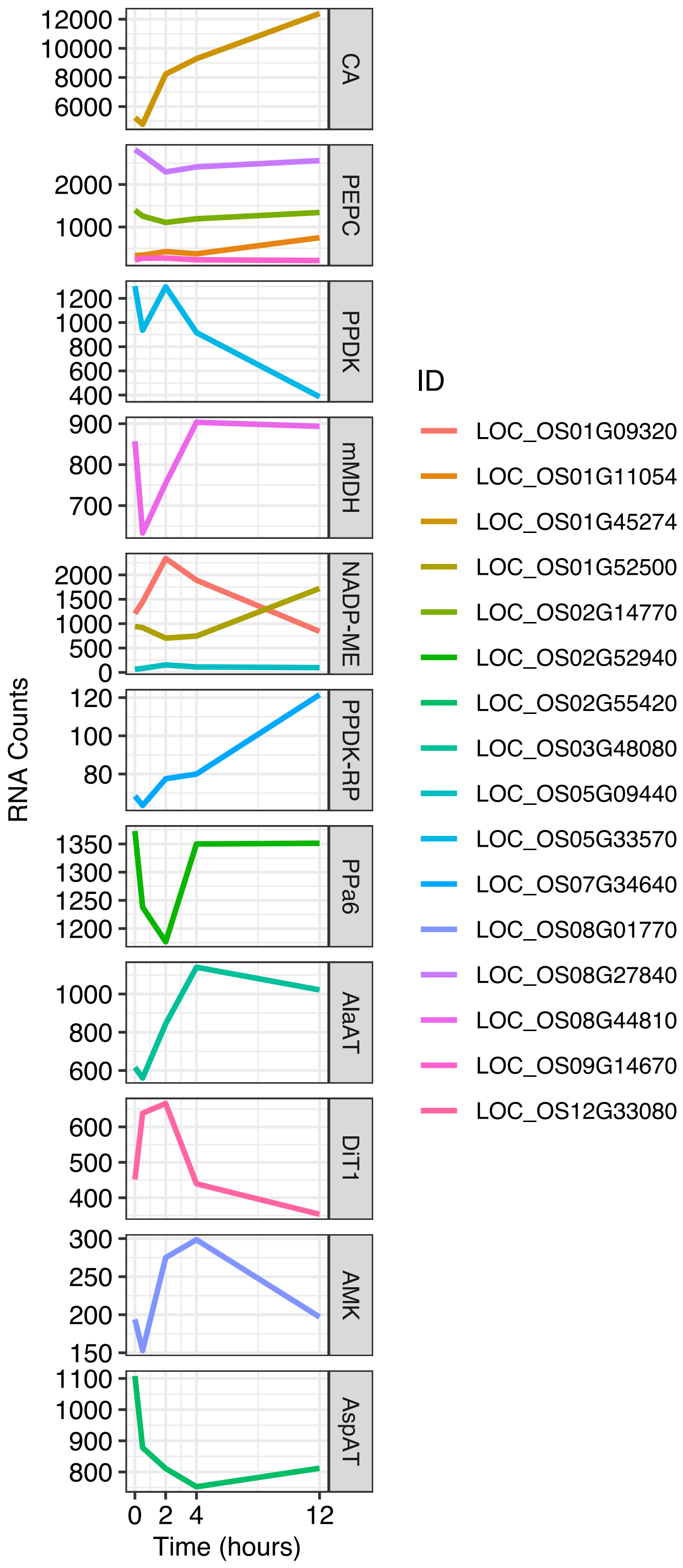

RNA Counts
Time (hours)
0
2
4
12
CA
PEPC
PPDK
mMDH
NADP-ME
PPDK-RP
PPa6
AlaAT
DiT1
AMK
AspAT
ID
LOC_OS01G09320
LOC_OS01G11054
LOC_OS01G45274
LOC_OS01G52500
LOC_OS02G14770
LOC_OS02G52940
LOC_OS02G55420
LOC_OS03G48080
LOC_OS05G09440
LOC_OS05G33570
LOC_OS07G34640
LOC_OS08G01770
LOC_OS08G27840
LOC_OS08G44810
LOC_OS09G14670
LOC_OS12G33080


Figure S2

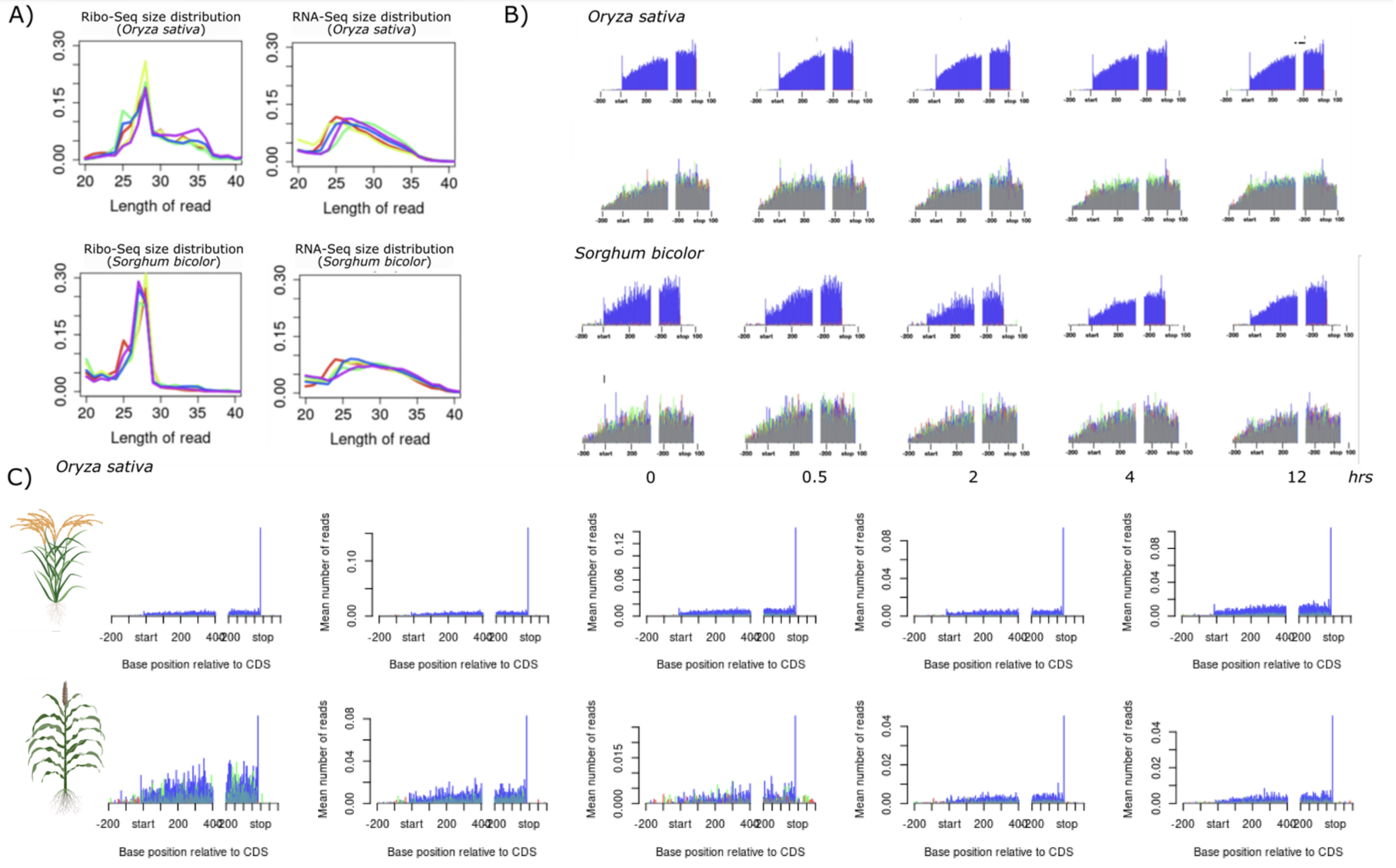
A)
Ribo-Seq size distribution (Oryza sativa)
RNA-Seq size distribution (Oryza sativa)
Ribo-Seq size distribution (Sorghum bicolor)
RNA-Seq size distribution (Sorghum bicolor)
Length of read
B)
Oryza sativa
Sorghum bicolor
0
0.5
2
4
12
hrs
C)
Oryza sativa
Mean number of reads
Base position relative to CDS
-200
start
200
400
stop

Figure S3

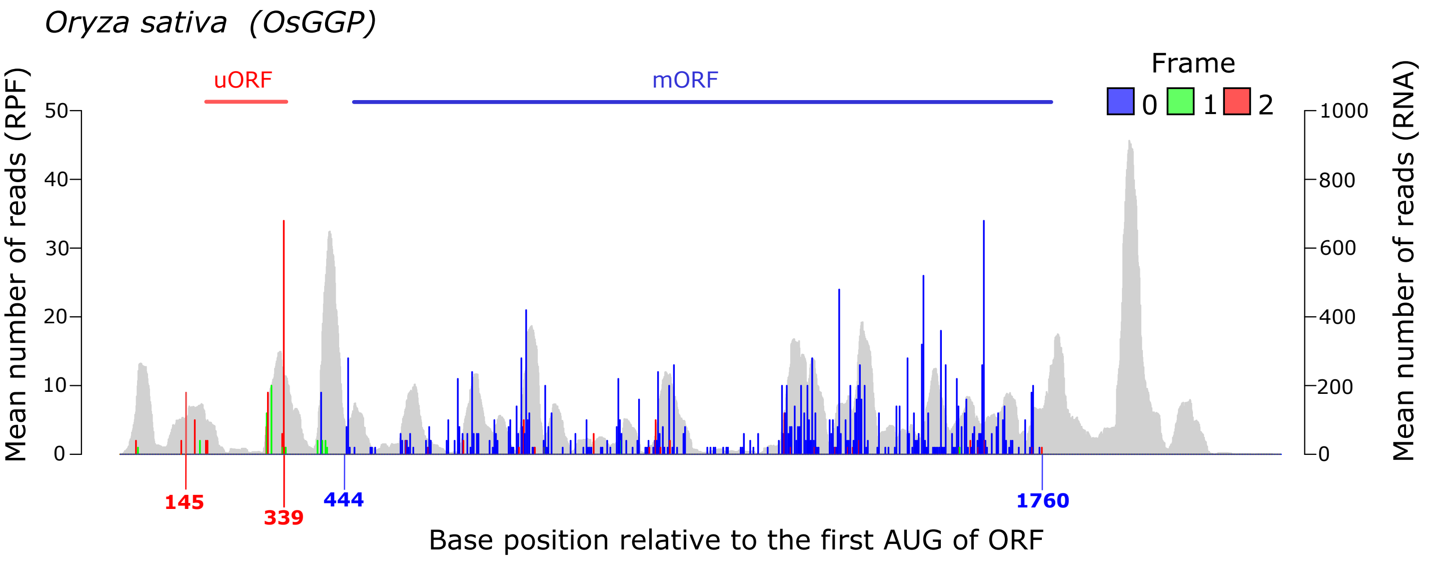

Oryza sativa (OsGGP)
uORF
mORF
Frame
0 1 2
Mean number of reads (RPF)
Mean number of reads (RNA)
50
40
30
20
10
0
1000
800
600
400
200
0
145
339
444
1760
Base position relative to the first AUG of ORF


Figure S4

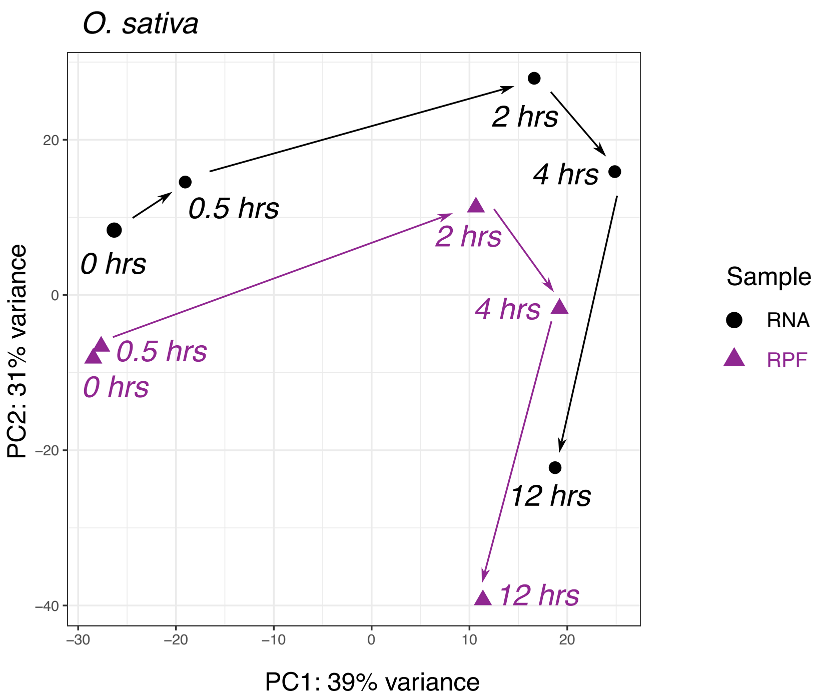

O. sativa
PC2: 31% variance
PC1: 39% variance
20
0
−20
−40
−30
−20
−10
0
10
20
0 hrs
0.5 hrs
2 hrs
4 hrs
12 hrs
0.5 hrs
0 hrs
2 hrs
4 hrs
12 hrs
Sample
RNA
RPF


### Figure S5

*O. sativa*

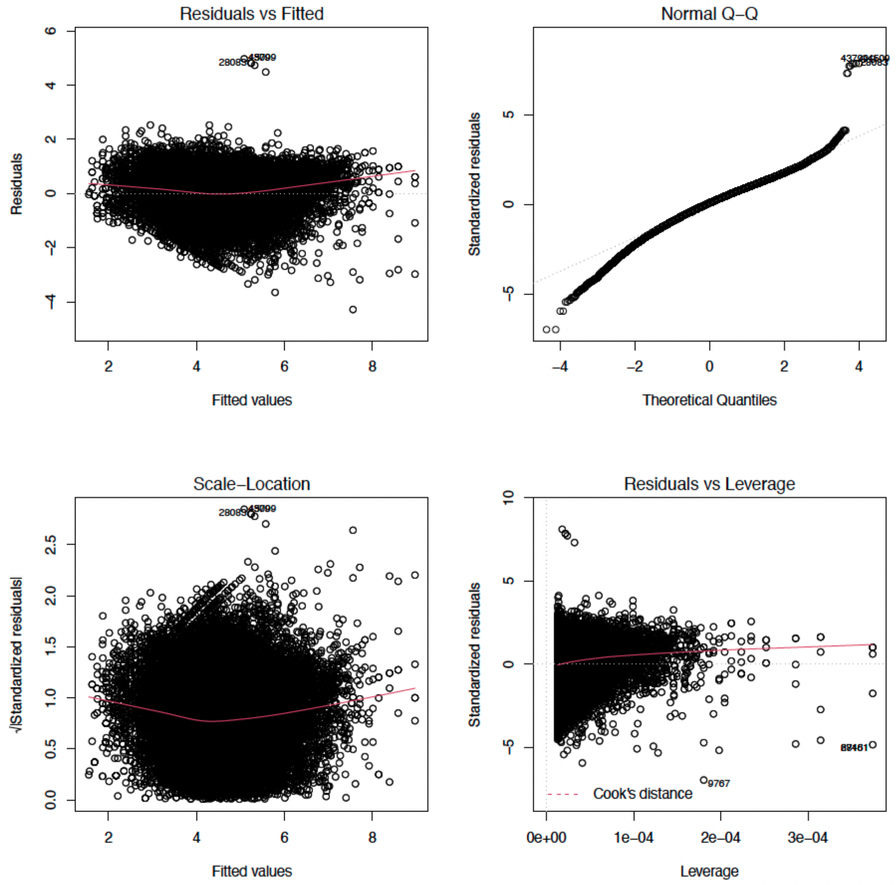


### Figure S6

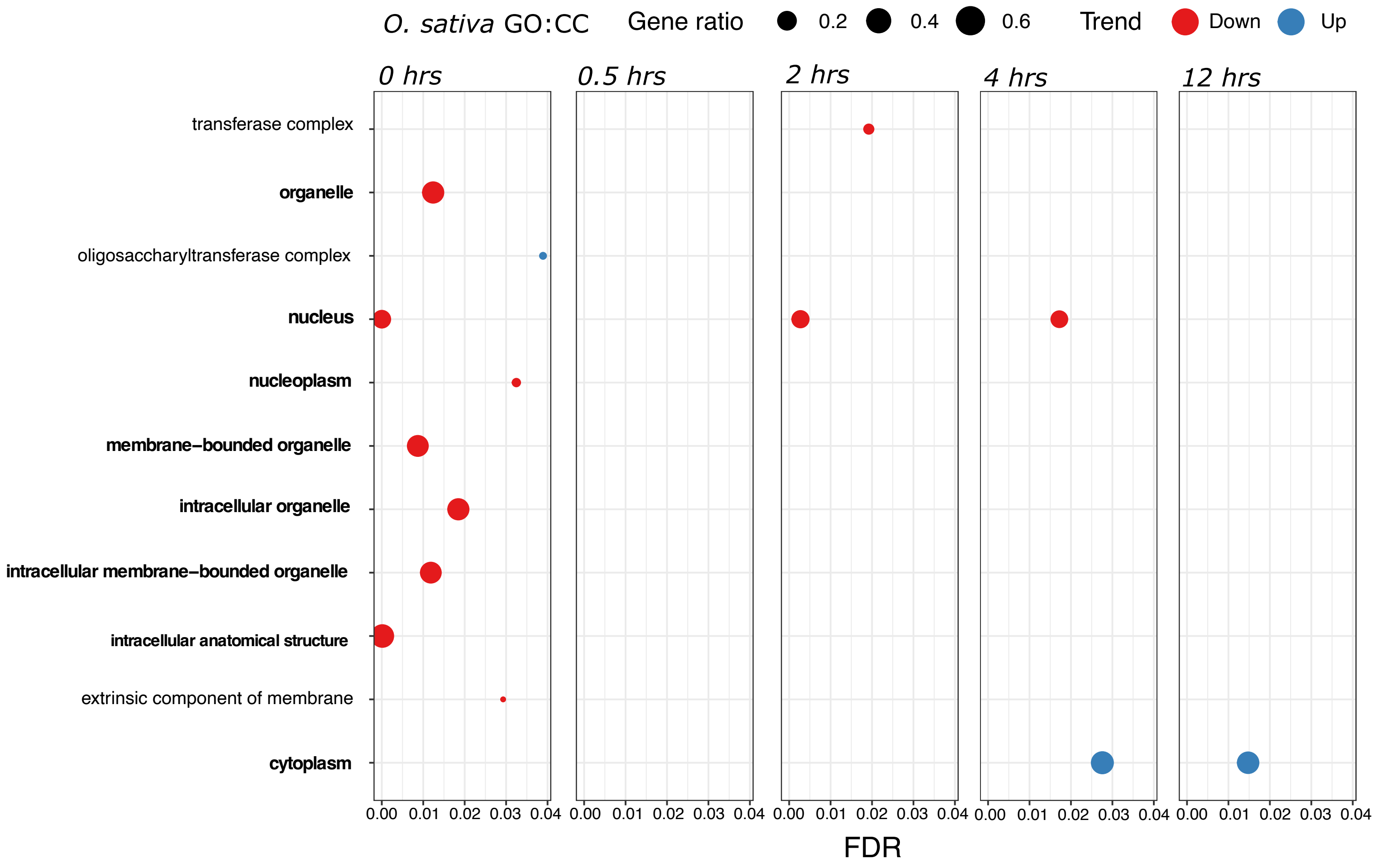
O. sativa GO:CC
Gene ratio
0.2
0.4
0.6
Trend
Down
Up
0 hrs
0.5 hrs
2 hrs
4 hrs
12 hrs
transferase complex
organelle
oligosaccharyltransferase complex
nucleus
nucleoplasm
membrane–bounded organelle
intracellular organelle
intracellular membrane–bounded organelle
intracellular anatomical structure
extrinsic component of membrane
cytoplasm
0.00 0.01 0.02 0.03 0.04
FDR

### Figure S7

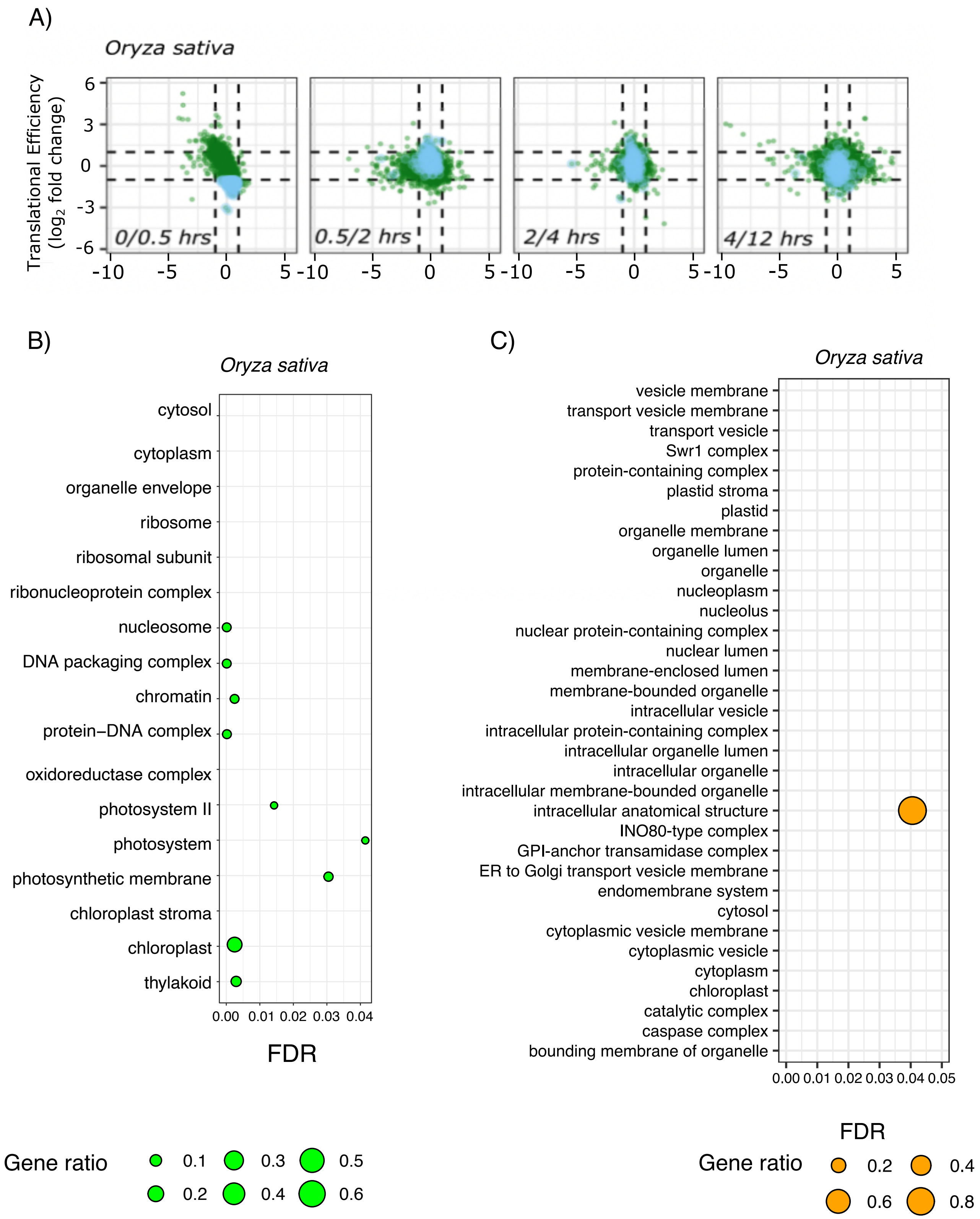
A)
Oryza sativa
Translational Efficiency (log2 fold change)
0/0.5 hrs
0.5/2 hrs
2/4 hrs
4/12 hrs
B)
Oryza sativa
cytosol
cytoplasm
organelle envelope
ribosome
ribosomal subunit
ribonucleoprotein complex
nucleosome
DNA packaging complex
chromatin
protein–DNA complex
oxidoreductase complex
photosystem II
photosystem
photosynthetic membrane
chloroplast stroma
chloroplast
thylakoid
FDR
Gene ratio
0.1 0.2 0.3 0.4 0.5 0.6
C)
Oryza sativa
vesicle membrane
transport vesicle membrane
transport vesicle
Swr1 complex
protein-containing complex
plastid stroma
plastid
organelle membrane
organelle lumen
organelle
nucleoplasm
nucleolus
nuclear protein-containing complex
nuclear lumen
membrane-enclosed lumen
membrane-bounded organelle
intracellular vesicle
intracellular protein-containing complex
intracellular organelle lumen
intracellular organelle
intracellular membrane-bounded organelle
intracellular anatomical structure
INO80-type complex
GPI-anchor transamidase complex
ER to Golgi transport vesicle membrane
endomembrane system
cytosol
cytoplasmic vesicle membrane
cytoplasmic vesicle
cytoplasm
chloroplast
catalytic complex
caspase complex
bounding membrane of organelle
FDR
Gene ratio
0.2 0.4 0.6 0.8

### Figure S8

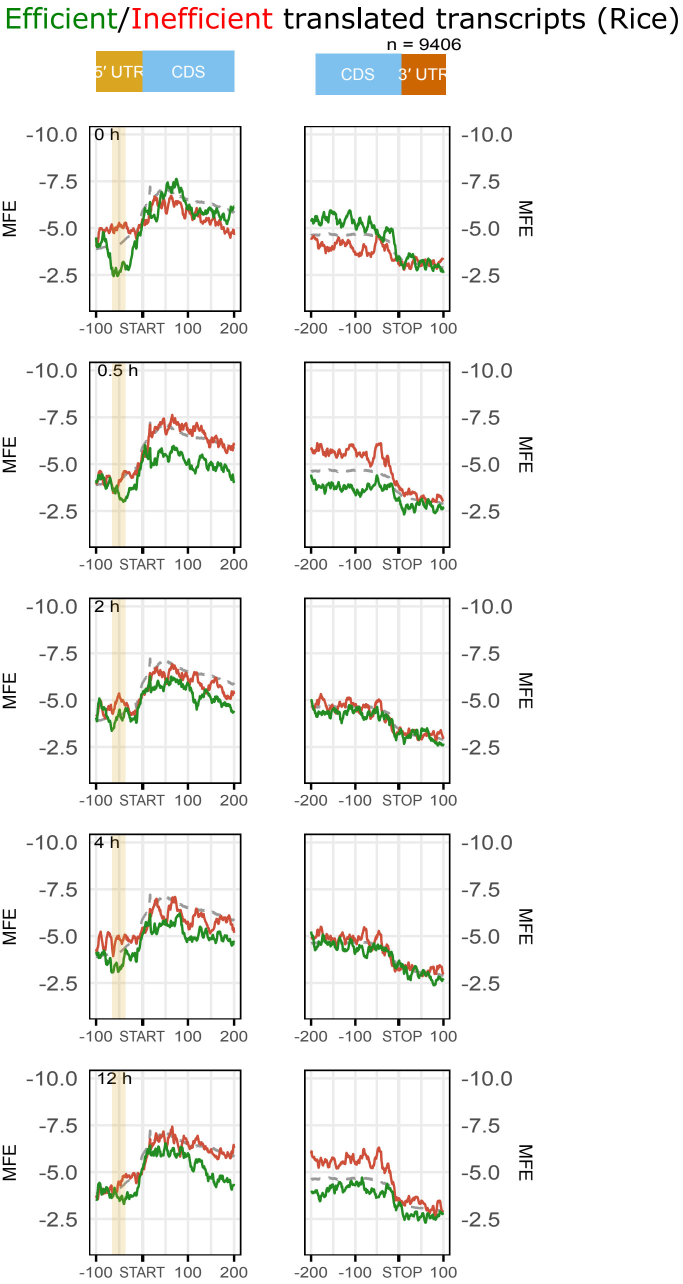
Efficient/Inefficient translated transcripts (Rice)
n = 9406
5′ UTR
CDS
CDS
3′ UTR
0 h
0.5 h
2 h
4 h
12 h
MFE
-10.0
-7.5
-5.0
-2.5
-100
START
100
200
-200
-100
STOP
100

### Figure S9

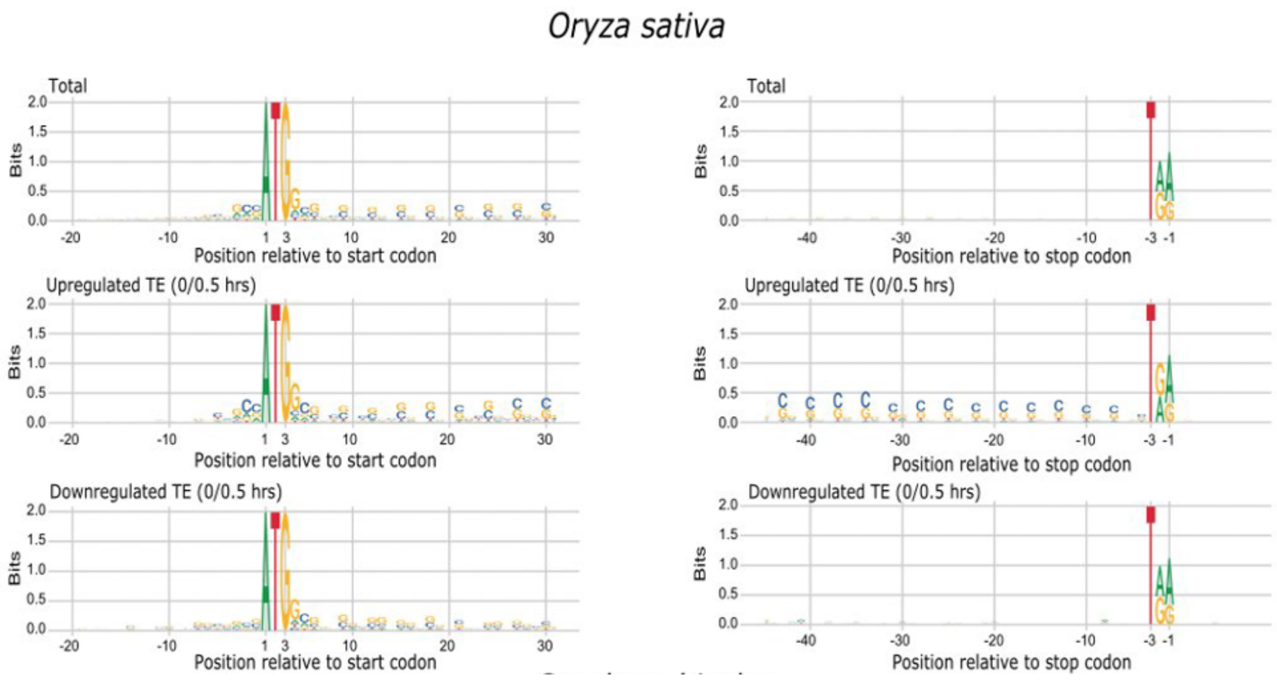
Oryza sativa
Total
Upregulated TE (0/0.5 hrs)
Downregulated TE (0/0.5 hrs)
Bits
2.0
1.5
1.0
0.5
0.0
-20
-10
1
3
10
20
30
Position relative to start codon
-40
-30
-20
-10
-3
-1
Position relative to stop codon

Figure S10

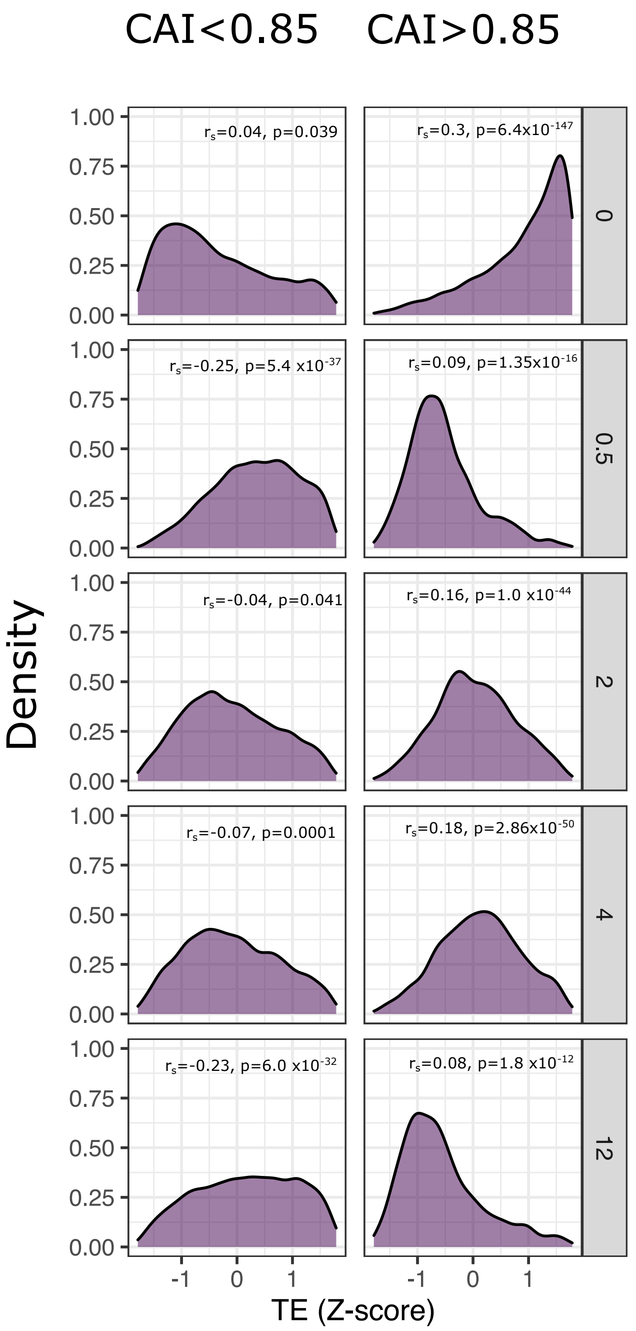
CAI<0.85
CAI>0.85
Density
TE (Z-score)
0
0.5
2
4
12
rs=0.04, p=0.039
rs=0.3, p=6.4x10-147
rs=-0.25, p=5.4 x10-37
rs=0.09, p=1.35x10-16
rs=-0.04, p=0.041
rs=0.16, p=1.0 x10-44
rs=-0.07, p=0.0001
rs=0.18, p=2.86x10-50
rs=-0.23, p=6.0 x10-32
rs=0.08, p=1.8 x10-12

Figure S11

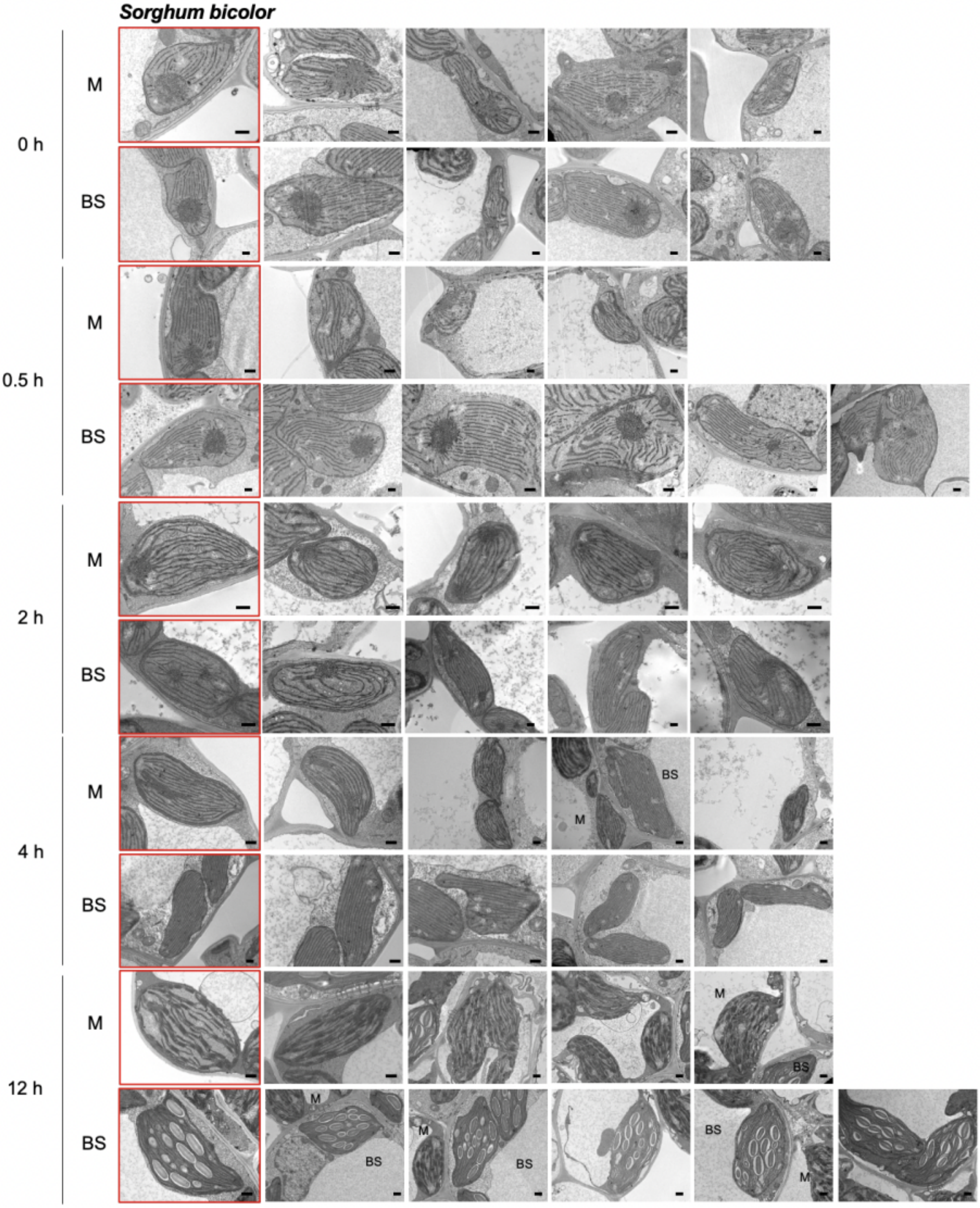


Scale bar = 500 nm

### Figure S12

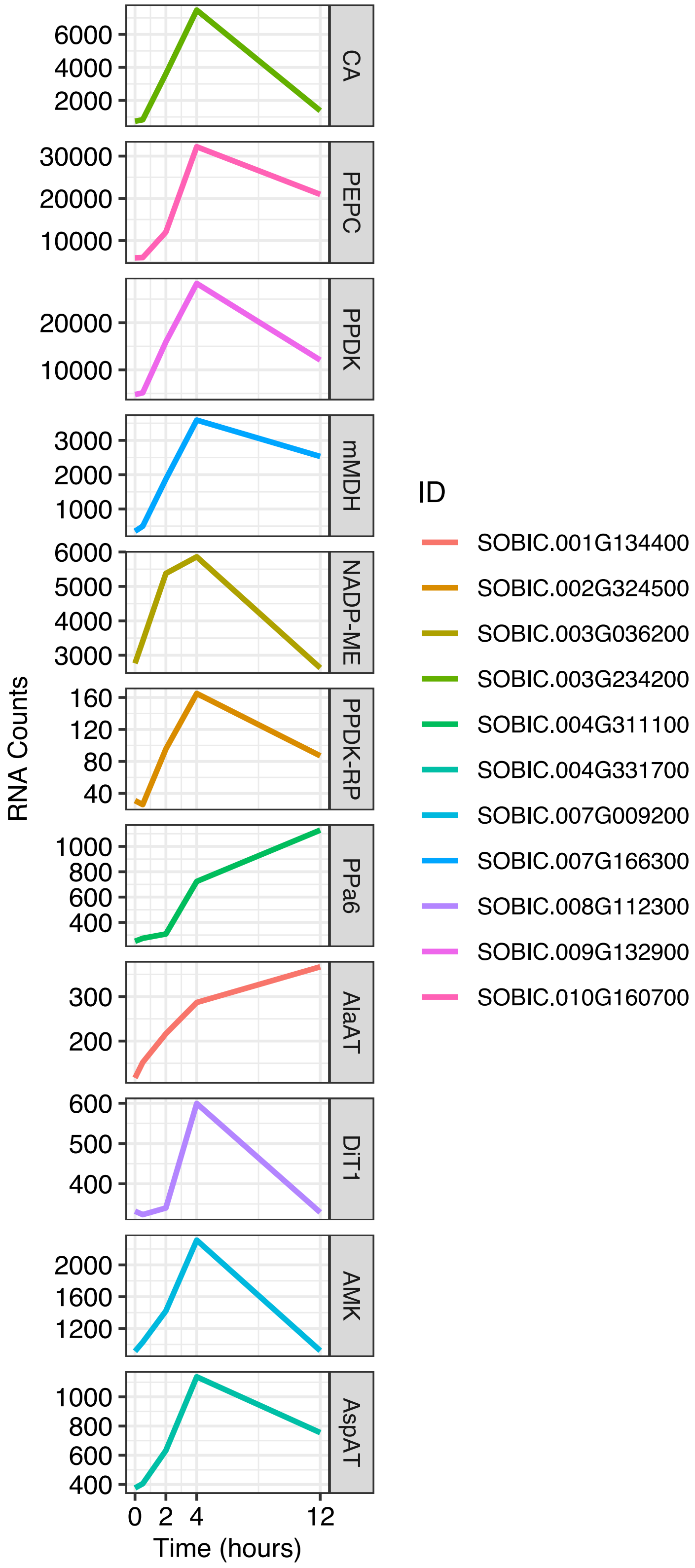
RNA Counts
CA
PEPC
PPDK
mMDH
NADP-ME
PPDK-RP
PPa6
AlaAT
DiT1
AMK
AspAT
Time (hours)
ID
SOBIC.001G134400
SOBIC.002G324500
SOBIC.003G036200
SOBIC.003G234200
SOBIC.004G311100
SOBIC.004G331700
SOBIC.007G009200
SOBIC.007G166300
SOBIC.008G112300
SOBIC.009G132900
SOBIC.010G160700

### Figure S13

*S. bicolor*

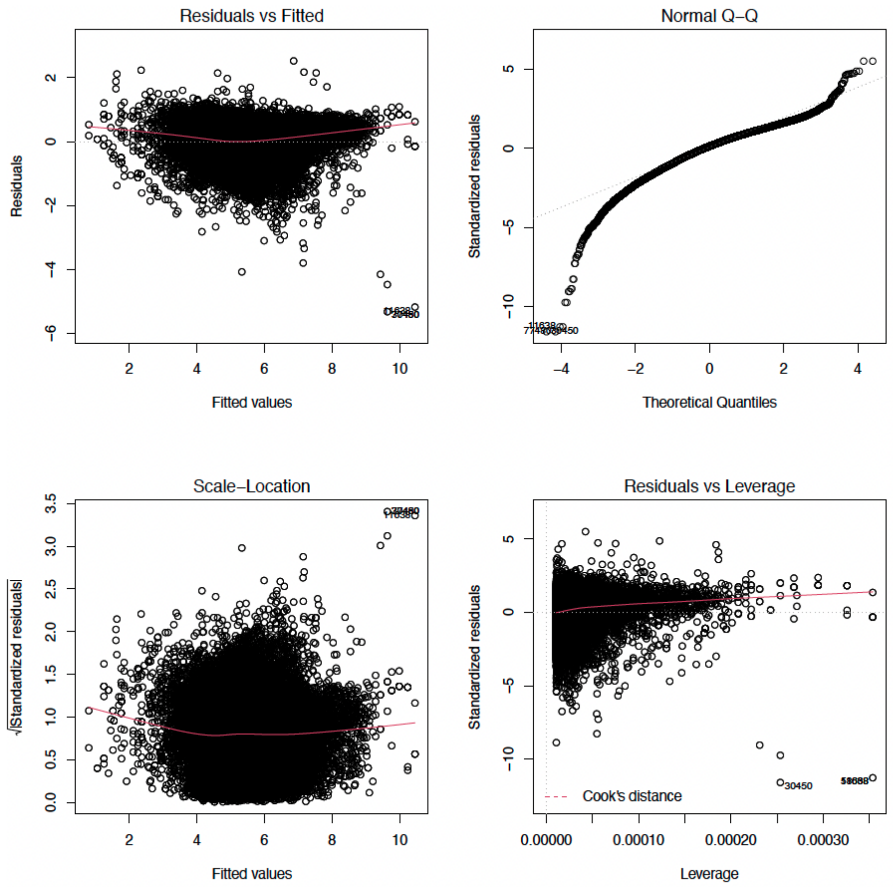


Figure S14

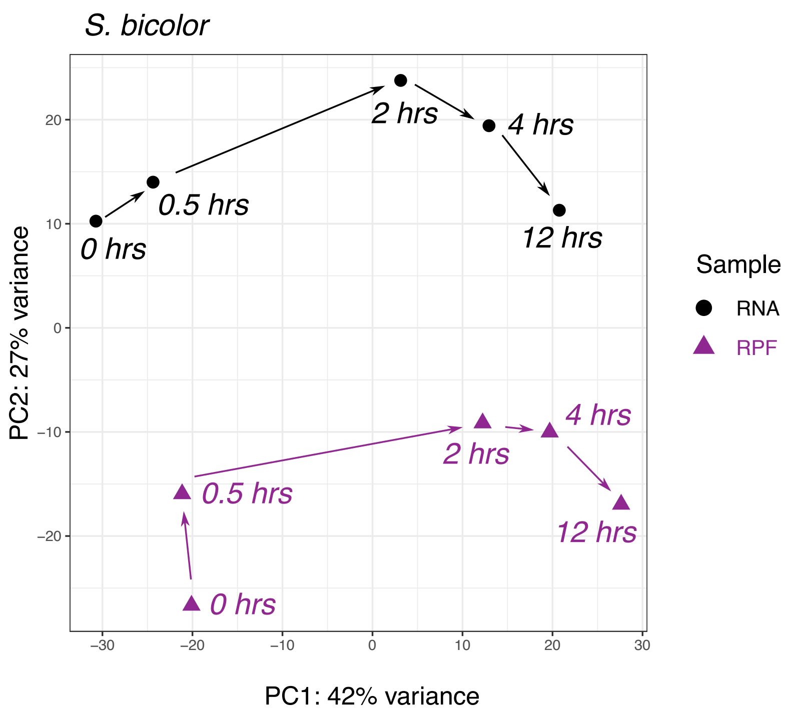

S. bicolor
2 hrs
4 hrs
0.5 hrs
0 hrs
12 hrs
4 hrs
2 hrs
0.5 hrs
12 hrs
0 hrs
20
10
0
−10
−20
−30
−20
−10
0
10
20
30
PC2: 27% variance
PC1: 42% variance
Sample
RNA
RPF


### Figure S15

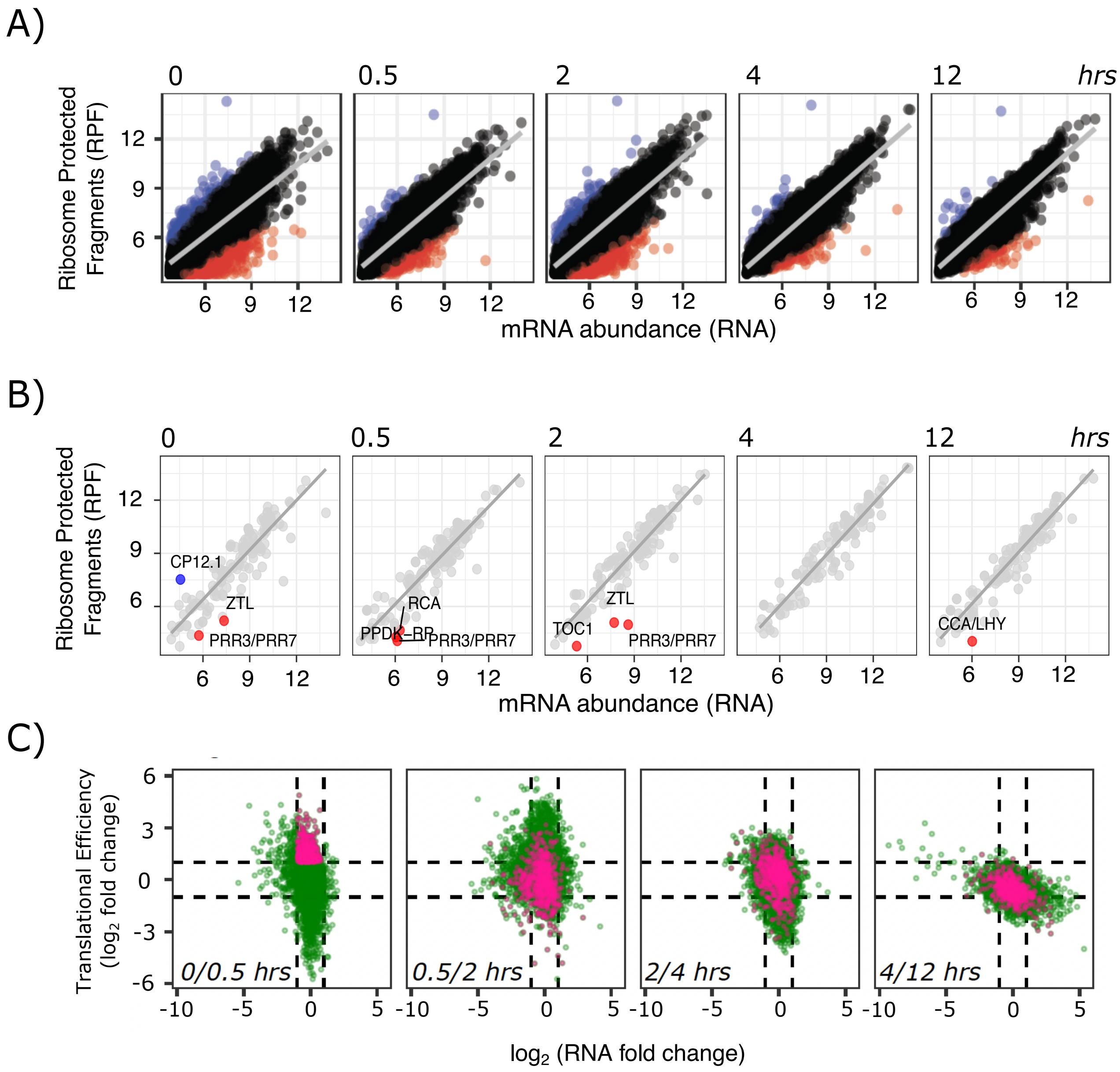
A)
0
0.5
2
4
12
hrs
Ribosome Protected Fragments (RPF)
mRNA abundance (RNA)
B)
CP12.1
ZTL
PRR3/PRR7
RCA
PPDK–RP
TOC1
CCA/LHY
C)
Translational Efficiency (log2 fold change)
0/0.5 hrs
0.5/2 hrs
2/4 hrs
4/12 hrs
log2 (RNA fold change)

### Figure S16

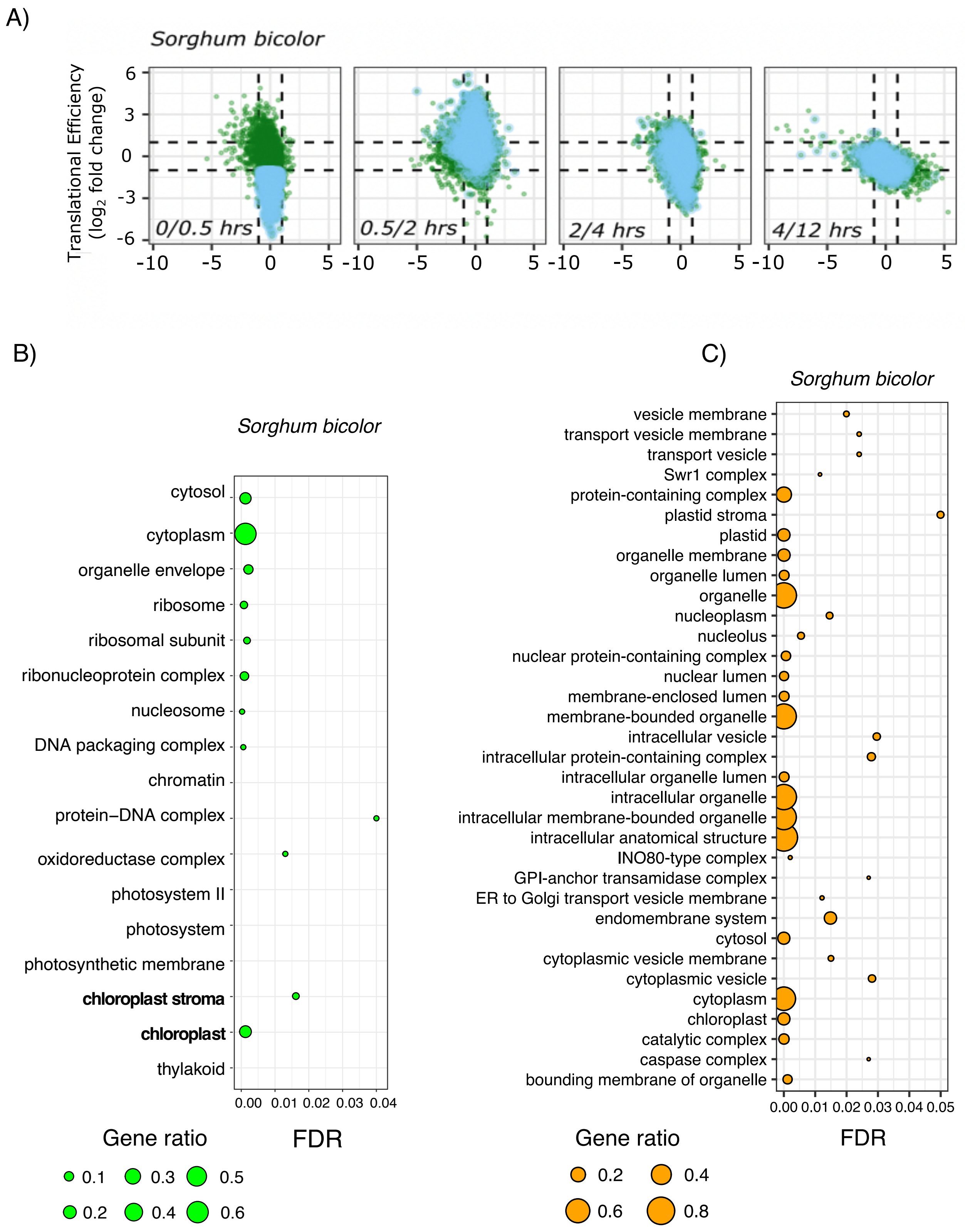
A)
Sorghum bicolor
Translational Efficiency
(log2 fold change)
6
3
0
-3
-6
0/0.5 hrs
0.5/2 hrs
2/4 hrs
4/12 hrs
-10 -5 0 5
B)
Sorghum bicolor
cytosol
cytoplasm
organelle envelope
ribosome
ribosomal subunit
ribonucleoprotein complex
nucleosome
DNA packaging complex
chromatin
protein–DNA complex
oxidoreductase complex
photosystem II
photosystem
photosynthetic membrane
chloroplast stroma
chloroplast
thylakoid
0.00 0.01 0.02 0.03 0.04
Gene ratio
FDR
0.1 0.3 0.5
0.2 0.4 0.6
C)
Sorghum bicolor
vesicle membrane
transport vesicle membrane
transport vesicle
Swr1 complex
protein-containing complex
plastid stroma
plastid
organelle membrane
organelle lumen
organelle
nucleoplasm
nucleolus
nuclear protein-containing complex
nuclear lumen
membrane-enclosed lumen
membrane-bounded organelle
intracellular vesicle
intracellular protein-containing complex
intracellular organelle lumen
intracellular organelle
intracellular membrane-bounded organelle
intracellular anatomical structure
INO80-type complex
GPI-anchor transamidase complex
ER to Golgi transport vesicle membrane
endomembrane system
cytosol
cytoplasmic vesicle membrane
cytoplasmic vesicle
cytoplasm
chloroplast
catalytic complex
caspase complex
bounding membrane of organelle
0.00 0.01 0.02 0.03 0.04 0.05
Gene ratio
FDR
0.2 0.4
0.6 0.8

### Figure S17

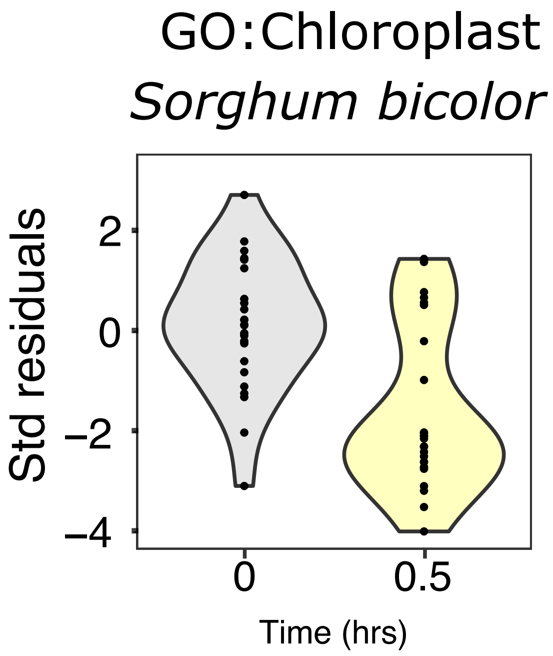

GO:Chloroplast
Sorghum bicolor
Std residuals
2
0
−2
−4
0
0.5
Time (hrs)


Figure S18

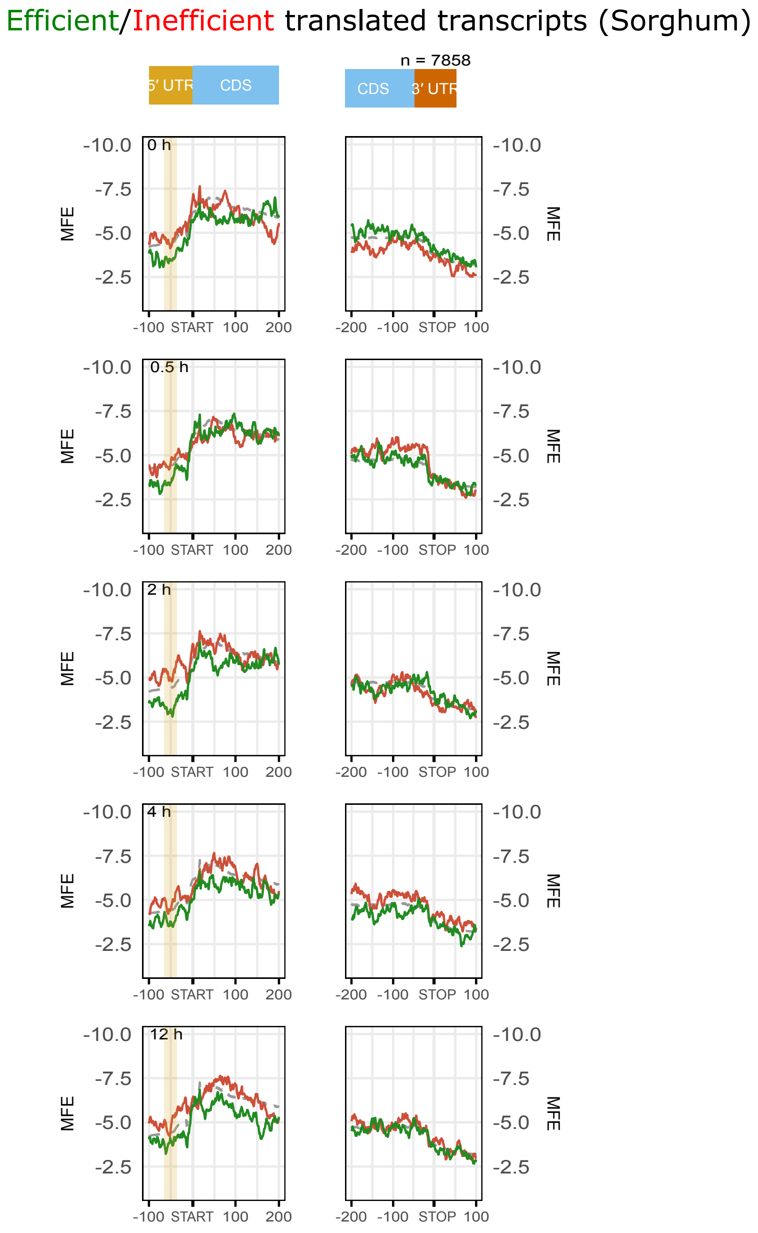

Efficient/Inefficient translated transcripts (Sorghum)
n = 7858
5′ UTR
CDS
CDS
3′ UTR
0 h
0.5 h
2 h
4 h
12 h
MFE
-10.0
-7.5
-5.0
-2.5
-100
START
100
200
-200
-100
STOP
100


### Figure S19

Figure S20

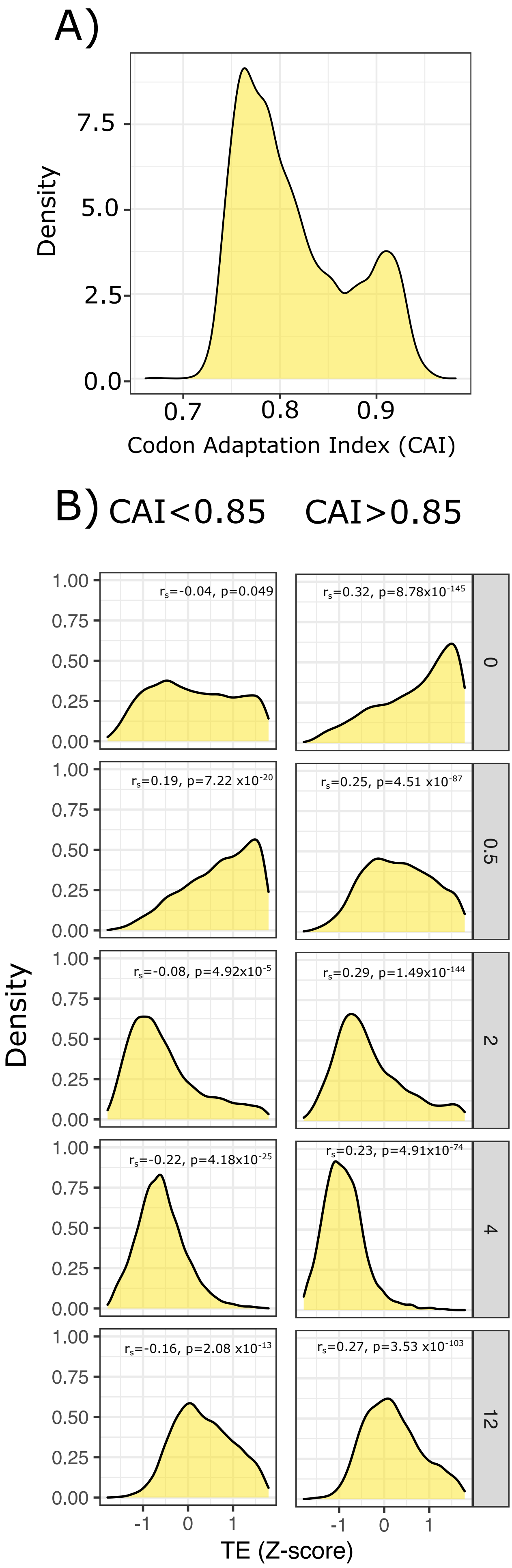
A)
Density
Codon Adaptation Index (CAI)
B)
CAI<0.85
CAI>0.85
$r_s$=-0.04, p=0.049
$r_s$=0.32, p=8.78x$10^{-145}$
$r_s$=0.19, p=7.22 x$10^{-20}$
$r_s$=0.25, p=4.51 x$10^{-87}$
$r_s$=-0.08, p=4.92x$10^{-5}$
$r_s$=0.29, p=1.49x$10^{-144}$
$r_s$=-0.22, p=4.18x$10^{-25}$
$r_s$=0.23, p=4.91x$10^{-74}$
$r_s$=-0.16, p=2.08 x$10^{-13}$
$r_s$=0.27, p=3.53 x$10^{-103}$
0
0.5
2
4
12
Density
TE (Z-score)

### Figure S21

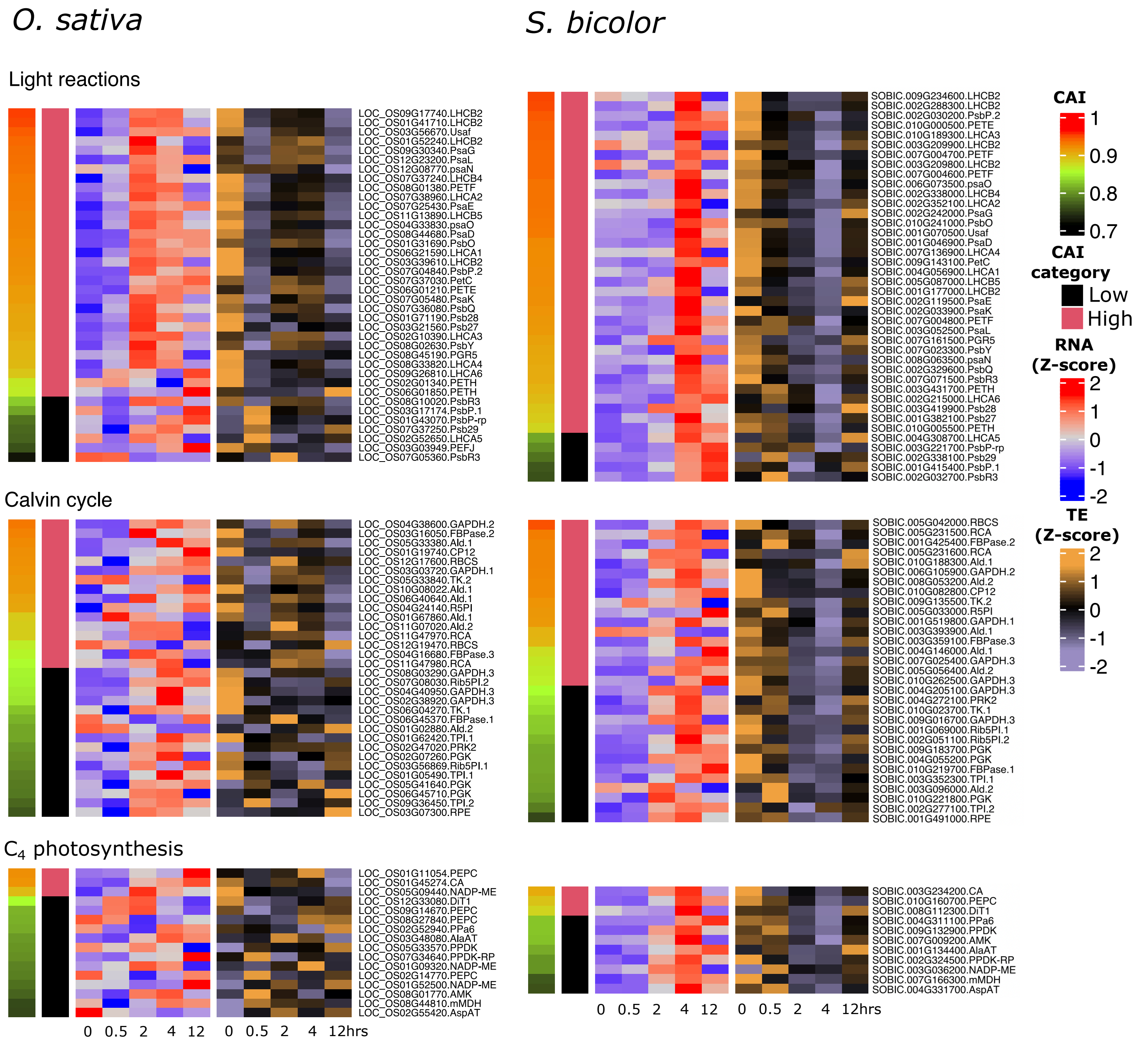

O. sativa
S. bicolor
Light reactions
LOC_OS09G17740.LHCB2
LOC_OS01G41710.LHCB2
LOC_OS03G56670.Usaf
LOC_OS01G52240.LHCB2
LOC_OS09G30340.PsaG
LOC_OS12G23200.PsaL
LOC_OS12G08770.psaN
LOC_OS07G37240.LHCB4
LOC_OS08G01380.PETF
LOC_OS07G38960.LHCA2
LOC_OS07G25430.PsaE
LOC_OS11G13890.LHCB5
LOC_OS04G33830.psaO
LOC_OS08G44680.PsaD
LOC_OS01G31690.PsbO
LOC_OS06G21590.LHCA1
LOC_OS03G39610.LHCB2
LOC_OS07G04840.PsbP.2
LOC_OS07G37030.PetC
LOC_OS06G01210.PETE
LOC_OS07G05480.PsaK
LOC_OS07G36080.PsbQ
LOC_OS01G71190.Psb28
LOC_OS03G21560.Psb27
LOC_OS02G10390.LHCA3
LOC_OS08G02630.PsbY
LOC_OS08G45190.PGR5
LOC_OS08G33820.LHCA4
LOC_OS09G26810.LHCA6
LOC_OS02G01340.PETH
LOC_OS01G01850.PETH
LOC_OS08G10020.PsbR3
LOC_OS03G17174.PsbP.1
LOC_OS01G43070.PsbP-rp
LOC_OS07G37250.Psb29
LOC_OS02G52650.LHCA5
LOC_OS03G03949.PEFJ
LOC_OS07G05360.PsbR3
SOBIC.009G234600.LHCB2
SOBIC.002G288300.LHCB2
SOBIC.002G030200.PsbP.2
SOBIC.010G000500.PETE
SOBIC.010G189300.LHCA3
SOBIC.003G209900.LHCB2
SOBIC.007G004700.PETF
SOBIC.003G209800.LHCB2
SOBIC.007G004600.PETF
SOBIC.006G073500.psaO
SOBIC.002G338000.LHCB4
SOBIC.002G352100.LHCA2
SOBIC.002G242000.PsaG
SOBIC.010G241000.PsbO
SOBIC.001G070500.Usaf
SOBIC.001G046900.PsaD
SOBIC.007G136900.LHCA4
SOBIC.009G143100.PetC
SOBIC.004G056900.LHCA1
SOBIC.005G087000.LHCB5
SOBIC.001G177000.LHCB2
SOBIC.002G119500.PsaE
SOBIC.002G033900.PsaK
SOBIC.007G004800.PETF
SOBIC.003G052500.PsaL
SOBIC.007G161500.PGR5
SOBIC.007G023300.PsbY
SOBIC.008G063500.psaN
SOBIC.002G329600.PsbQ
SOBIC.007G071500.PsbR3
SOBIC.003G431700.PETH
SOBIC.002G215000.LHCA6
SOBIC.003G419900.Psb28
SOBIC.001G382100.Psb27
SOBIC.010G005500.PETH
SOBIC.004G308700.LHCA5
SOBIC.003G221700.PsbP-rp
SOBIC.002G338100.Psb29
SOBIC.001G415400.PsbP.1
SOBIC.002G032700.PsbR3
CAI
1
0.9
0.8
0.7
CAI category
Low
High
RNA (Z-score)
2
1
0
-1
-2
TE (Z-score)
2
1
0
-1
-2
Calvin cycle
LOC_OS04G38600.GAPDH.2
LOC_OS03G16050.FBPase.2
LOC_OS05G33380.Ald.1
LOC_OS01G19740.CP12
LOC_OS12G17600.RBCS
LOC_OS03G03720.GAPDH.1
LOC_OS05G33840.TK.2
LOC_OS10G08022.Ald.1
LOC_OS06G40640.Ald.1
LOC_OS04G24140.R5PI
LOC_OS01G67860.Ald.1
LOC_OS11G07020.Ald.2
LOC_OS11G47970.RCA
LOC_OS12G19470.RBCS
LOC_OS04G16680.FBPase.3
LOC_OS11G47980.RCA
LOC_OS08G03290.GAPDH.3
LOC_OS07G08030.Rib5PI.2
LOC_OS04G40950.GAPDH.3
LOC_OS02G38920.GAPDH.3
LOC_OS06G04270.TK.1
LOC_OS06G45370.FBPase.1
LOC_OS01G02880.Ald.2
LOC_OS01G62420.TPI.1
LOC_OS02G47020.PRK2
LOC_OS02G07260.PGK
LOC_OS03G56869.Rib5PI.1
LOC_OS01G05490.TPI.1
LOC_OS05G41640.PGK
LOC_OS06G45710.PGK
LOC_OS09G36450.TPI.2
LOC_OS03G07300.RPE
SOBIC.005G042000.RBCS
SOBIC.005G231500.RCA
SOBIC.001G425400.FBPase.2
SOBIC.005G231600.RCA
SOBIC.010G188300.Ald.1
SOBIC.006G105900.GAPDH.2
SOBIC.008G053200.Ald.2
SOBIC.010G082800.CP12
SOBIC.009G135500.TK.2
SOBIC.005G033000.R5PI
SOBIC.001G519800.GAPDH.1
SOBIC.003G393900.Ald.1
SOBIC.003G359100.FBPase.3
SOBIC.004G146000.Ald.1
SOBIC.007G025400.GAPDH.3
SOBIC.005G056400.Ald.2
SOBIC.010G262500.GAPDH.3
SOBIC.004G205100.GAPDH.3
SOBIC.004G272100.PRK2
SOBIC.010G023700.TK.1
SOBIC.009G016700.GAPDH.3
SOBIC.001G069000.Rib5PI.1
SOBIC.002G051100.Rib5PI.2
SOBIC.009G183700.PGK
SOBIC.004G055200.PGK
SOBIC.010G219700.FBPase.1
SOBIC.003G352300.TPI.1
SOBIC.003G096000.Ald.2
SOBIC.010G221800.PGK
SOBIC.002G277100.TPI.2
SOBIC.001G491000.RPE
C4 photosynthesis
LOC_OS01G11054.PEPC
LOC_OS01G45274.CA
LOC_OS05G09440.NADP-ME
LOC_OS12G33080.DiT1
LOC_OS09G14670.PEPC
LOC_OS08G27840.PEPC
LOC_OS02G52940.PPa6
LOC_OS03G48080.AlaAT
LOC_OS05G33570.PPDK
LOC_OS07G34640.PPDK-RP
LOC_OS01G09320.NADP-ME
LOC_OS02G14770.PEPC
LOC_OS01G52500.NADP-ME
LOC_OS08G01770.AMK
LOC_OS08G44810.mMDH
LOC_OS02G55420.AspAT
SOBIC.003G234200.CA
SOBIC.010G160700.PEPC
SOBIC.008G112300.DiT1
SOBIC.004G311100.PPa6
SOBIC.009G132900.PPDK
SOBIC.007G009200.AMK
SOBIC.001G134400.AlaAT
SOBIC.002G324500.PPDK-RP
SOBIC.003G036200.NADP-ME
SOBIC.007G166300.mMDH
SOBIC.004G331700.AspAT
0 0.5 2 4 12 0 0.5 2 4 12hrs
0 0.5 2 4 12 0 0.5 2 4 12hrs


### Figure S22

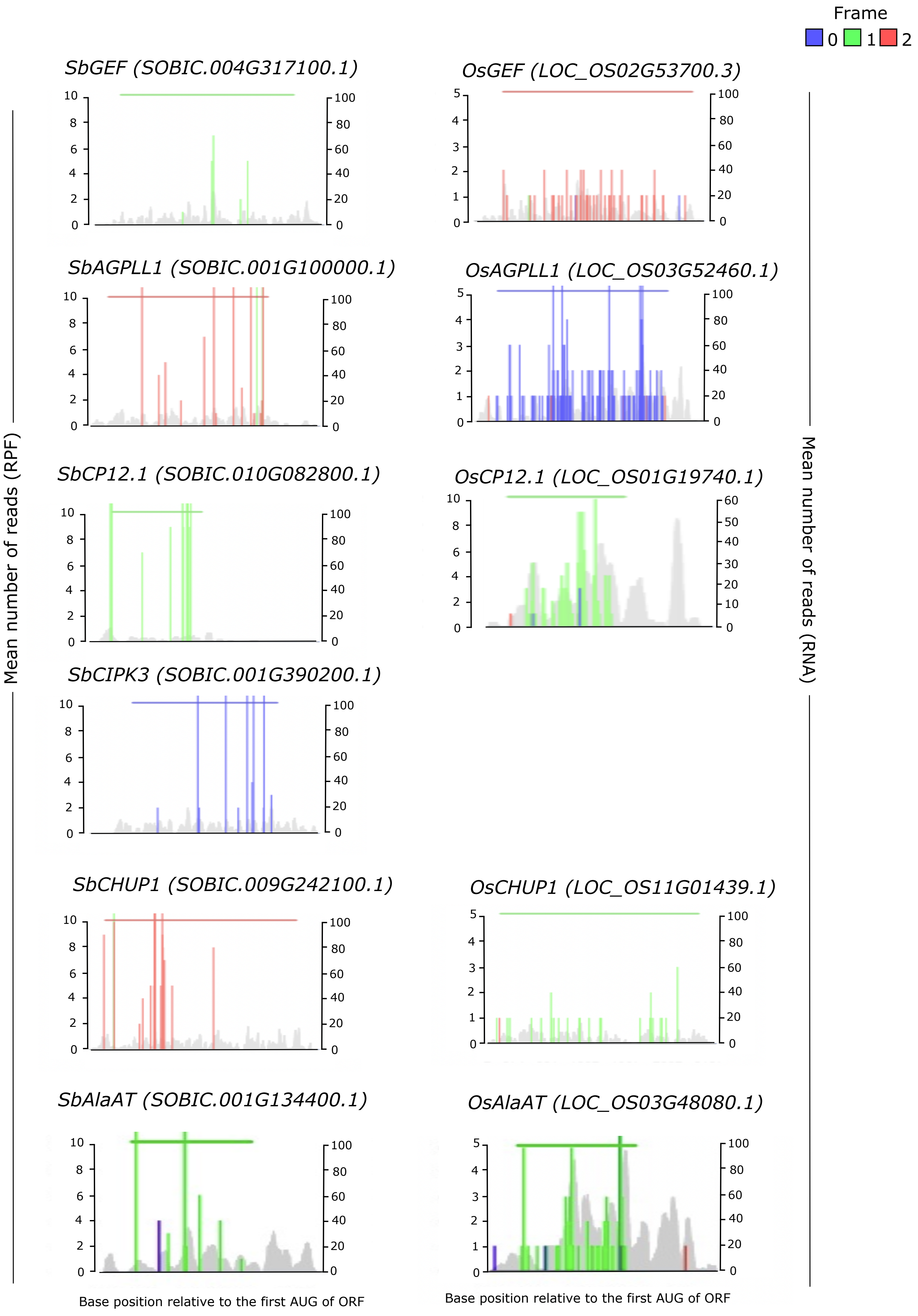
Frame
0 1 2
SbGEF (SOBIC.004G317100.1)
OsGEF (LOC_OS02G53700.3)
SbAGPLL1 (SOBIC.001G100000.1)
OsAGPLL1 (LOC_OS03G52460.1)
SbCP12.1 (SOBIC.010G082800.1)
OsCP12.1 (LOC_OS01G19740.1)
SbCIPK3 (SOBIC.001G390200.1)
SbCHUP1 (SOBIC.009G242100.1)
OsCHUP1 (LOC_OS11G01439.1)
SbAlaAT (SOBIC.001G134400.1)
OsAlaAT (LOC_OS03G48080.1)
Mean number of reads (RPF)
Mean number of reads (RNA)
Base position relative to the first AUG of ORF
Base position relative to the first AUG of ORF

Figure S23
